# A novel GLYCEROPHOSPHODIESTER PHOSPHODIESTERASE 13 is involved in the Phosphate starvation-induced lipid remodeling in rice

**DOI:** 10.1101/2024.03.27.586571

**Authors:** Duong Thi Thuy Dang, Thi Linh Nguyen, Anh-Tuan Tran, Tam Thi Thanh Tran, Tuan-Anh Tran, Kieu Thi Xuan Vo, Jong-Seong Jeon, Tien Van Vu, Jae-Yean Kim, Phuong Nhue Nguyen, Hoang Ha Chu, Phat Tien Do, Huong Thi Mai To

## Abstract

Phosphorus (P) is one of the most vital macronutrient determinants in plant development and productivity. However, the bioavailability of inorganic phosphate (Pi), the only form that plants can assimilate, is limited in the soil, thus significantly affecting plant development. Plants have adopted various specialized strategies to modify their morphological, physiological, and biochemical properties for better adaptation to Pi deficiency conditions. GLYCEROPHOSPHODIESTER PHOSPHODIESTERASES (GDPDs), generally known as phospholipid remodeling proteins, have been suggested to play essential roles in maintaining phosphate homeostasis. The previous genome-wide association studies (GWAS) in a Vietnamese rice collection led to the discovery of a robust QTL named *qRST9.14* associating with the phosphate adaption in rice, in which *OsGDPD13* is located within this locus. Interestingly, we discovered an absence of *OsGDPD13* from the *Indica* reference genome. A subcellular localization study showed that the GDPD13 protein localizes to the plasma membrane, cytoplasmic speckles, and plasmodesmata. The *osgdpd13* knockout mutant was generated in the *Japonica* cv Kitaake background to characterize its function. Phenotypic analysis indicated that mutation of *OsGDPD13* significantly lowered crown root number and reduced the plant height under Pi deficient condition. Moreover, *osgdpd13* lines reduce the capability to degrade total phospholipids compared to the wild type under Pi starvation conditions. This finding suggests the involvement of the *OsGDPD13* gene in rice growth and the Phosphate starvation-induced lipid remodeling under P deficiency. This work will provide valuable information for developing crop plants with higher Phosphate use efficiency.

## INTRODUCTION

Rice (*Oryza sativa* L.) is vital in ensuring food security worldwide because it is consumed by more than half of the world’s population. Its importance is evident given the fact that rice cultivation accounts for nearly 20% of the world’s cereal production, and global rice consumption is expected to increase by 1.1% per annum (OECD/FAO, 2022). However, agricultural production faces numerous obstacles due to climate change, making food security challenging. Of the identified challenges, soil security is increasingly becoming widely discussed because sustainable soil function is essential for 95% of the global food supply (Pozza & Field, 2020). Soil fertility and nutrients, including macro- and micronutrients, are the mainstays of agriculture. The lack of soil nutrients causes defects in growth, reducing agricultural output and nutritional value (Vargas Rojas et al., 2022).

Phosphorus (P) is one of the most vital macronutrients for plant development, and productivity (Kumar et al., 2023; Siedliska et al., 2021). Numerous studies have shown that P is involved in a wide range of vital functions, including photosynthesis, seed germination, seedling development, root, shoot, flower, respiration, and nitrogen fixation (Malhotra et al., 2018) There are several chemical types of soil phosphorus, including inorganic phosphorus and organic phosphorus (Shen et al., 2011) and inorganic phosphate (Pi) is considered the only form that can be absorbed and assimilated by plants (Kumar et al., 2023). However, Pi has low availability in soil and poor accessibility to plants because it is easily fixed by soil mineral components, meaning Pi can be a hindering factor to plant growth (Ojeda-Rivera et al., 2022; Shen et al., 2011). This situation, coupled with the fact that P is a finite and non-renewable resource, facilitates the usage of chemical P fertilizers. The persistent over-application of P fertilizers has resulted in the accumulation of heavy metals in agricultural soil and eutrophication issues, which threaten the environment (De López Camelo et al., 1997). Hence, developing P-efficient crops has been one of the sustainable approaches to meet the growing demand for food while ensuring environmental health.

Developing crop varieties with low Pi tolerance and enhanced PUE is one of the critical tasks for researchers seeking to address the problem of Pi starvation (Yip et al., 2011). Significant efforts have been made worldwide to comprehend the molecular mechanisms of plants’ responses under Pi-limiting conditions to improve Pi utilization efficiency (PUE) in crops. Plants possess many sophisticated adaptive modifications, including morphological, biochemical, physiological, and molecular responses to better adapt to Pi-deficient conditions (Yuan & Liu, 2008). Plants could increase their Pi-acquisition efficiency (the ability of their roots to take up Pi for the soil) and/or PUE (Veneklaas et al., 2012). PUE is a complex quantitative trait associated with numerous genes; however, the underlying molecular mechanisms remain largely elusive (Zhang et al., 2014). Several genes or regulators primarily engaged in nutrient metabolism, absorption, or remobilization could be potential targets at the molecular level to increase plants’ PUE (Mehra & Giri, 2016). For instance, phospholipase C (PLC), phospholipase D (PLD), and Phosphatidic acid phosphohydrolase (PAH) increase the availability of P by releasing cellular phosphate from membrane phospholipids through phospholipid degradation (Nakamura, 2013). It has been demonstrated that plant phosphate transporters play a role in the uptake of Pi from the soil and its translocation in plants (Jia et al., 2011). The overexpression of the phosphate transporter gene *Pht1;1*, *Pht1;2*, *Pht1;3*, and *Pht1;4* in *Arabidopsis thaliana* (Mudge et al., 2002); *OsPht1;8* (Jia et al., 2011) and *OsPHT1;6* (Zhang et al., 2014) in rice resulted in greater Pi content under Pi starvation stress. In contrast, the knockdown of *OsPht1;8* reduced Pi uptake and plant growth under control and low Pi conditions (Jia et al., 2011).

Internal P remobilization is an essential rescue trait of plants to acclimate under P-deprived conditions (Hammond et al., 2004). Membrane lipid remodeling has drawn attention as a significant source of internal phosphate supply that plants can mobilize under phosphate deficiency conditions by releasing Pi from phospholipid head groups (Nakamura, 2013). Pi scarcity activates membrane lipid remodeling in higher plants, which appears to be an essential adaptation mechanism for plants in response to Pi starvation through the relocalization and recycling of internal Pi inside various cells (Liu et al., 2014). The major components of the P-free galactolipid in green tissue include 50–57% monogalactosyldiacylglycerol (MGDG), 15– 28% digalactosyldiacylglycerol (DGDG), 2–5% SQDG (Härtel et al., 2001). Most of the DGDG content is localized in plastidial membranes under Pi-sufficient conditions. Under Pi-deficient conditions, DGDG is transported to extra plastidial membranes to replace P-lipids (Jouhet et al., 2004). Glycerolipids sharing properties in common were considered to substitute for each other; for example, both DGDG and phosphatidylcholine (PC) consist of large polar head groups, are bilayer lipids, and may increase membrane stability. Glycerolipids of small head groups such as MGDG and phosphatidylethanolamine (PE) can form non-bilayer H_II_-type structures (Jouhet, 2013). In Pi-starved plants, Pi is released from membrane phospholipids by the hydrolyzation of the phosphate-containing headgroup and the replacement by the non-P-containing galactolipids for maintaining membrane integrity (Nakamura, 2013).

GLYCEROPHOSPHODIESTER PHOSPHODIESTERASE (GDPD) are enzymes involved in the degradation of glycerophosphodiesters into *sn*-glycerol-3-phosphate (G3P), which plays essential roles in maintaining phosphate homeostasis (Mehra & Giri, 2016). Under low Pi conditions, the expression levels of the GDPD genes are highly induced along with the increasing accumulation of Pi content, root growth, and biomass compared to wild-type, indicating their potential roles in the low P adaptation (Mehra et al., 2019). The induction of expression level was also observed in white lupin (Cheng et al. 2011a), *Arabidopsis* (Y. Cheng et al. 2011b), chickpea (Mehra & Giri, 2016), and rice (Mehra et al., 2019; Mehra & Giri, 2016). In *Arabidopsis*, *atgdpd1* mutants play a significant role in root formation under Pi-deficient conditions (Cheng et al. 2011b). The overexpression of *OsGDPD2* greatly enhanced the Pi content and biomass in rice under Pi starvation conditions, thereby successfully uncovering the significant contribution of this gene to rice’s low Pi tolerance (Mehra et al., 2019). Another *OsGDPD* gene, *OsGDPD5*, was strongly expressed, with its expression levels increasing by almost 147 times under low P (Mehra & Giri, 2016).

Two genome-wide association studies (GWAS) conducted with 160 indigenous Vietnamese rice genotypes have allowed us to identify some promising quantitative trait loci (QTLs) and candidate genes related to P efficiency (Mai et al., 2021; To et al., 2020). The most robust QTL discovered from these results is *qRST9.14,* located in Chromosome 9. The Linkage disequilibrium heatmap showed seven tightly linked markers inside the *qRST9.14* intervals, of which the marker *Dj09_10404317R* located in the intron acceptor splicing site between exon 3 and 4 of the *OsGDPD13* gene had the most significant p-value (*p* = 3.18e-10). Therefore, the *OsGDPD13* gene was functionally investigated to understand better its underlying mechanisms in plant adaptation to low Pi supply. The CRISPR-Cas9 system was used to generate *osgdpd13* mutants in *O. sativa* ssp. *japonica variety* Kitaake. T3 of two homozygous lines were selected for phenotyping rice’s physiological and biochemical parameters under the Pi starvation condition. Understanding how the *OsGDPD13* gene is involved in Pi starvation tolerance will provide valuable information for developing improved phosphate-efficient crops.

## MATERIALS AND METHODS

### Plant materials

*Oryza sativa* L. ssp *japonica* Kitaake mature seeds were used as the wild type for phenotyping and the genetic background for rice transformation and mutation. Four Vietnamese germplasms including G22, G99, G207 (*indica*), and G299 (*japonica*) were used for whole genome sequencing.

### Whole genome sequencing analysis

DNA samples were sent to sequencing using the Illumina platform (Novogene Biotech Co., Ltd, China) with 14x coverage or greater. The raw sequencing data were first preprocessed to eliminate poor-quality reads with Phred scores lower than 22 using Trimmomatic v.0.39 (Bolger et al., 2014). The processed reads were aligned to the *japonica* Nipponbare *OsGDPD13* genomic region and its promoter region (2000 bp long upstream of the Os*GDPD13* coding sequence), which were extracted from the Rice Genome Annotation Project database (http://rice.uga.edu/). These alignment processes utilized the Burrows-Wheeler Alignment (bwa) tool v.0.7.17 (Li and Durbin, 2009). Mapped reads from the two alignments were extracted using Samtools v.1.11 to construct contigs with SPAdes v.3.14.1 (Bankevich et al., 2012). The contigs below 200 bp were filtered out. The remaining contigs were then aligned back to the *japonica* reference sequence (GCA_001433935.1) using Minimap2 v2.24 (Li, 2018). These alignments were visualized on The Integrative Genomics Viewer (IGV) program version 2.26.1 (Robinson et al., 2011).

### *In silico* structure prediction and docking studies

*OsGDPD13* promoter region was queried for potential CAREs using the New PLACE database (https://www.dna.affrc.go.jp/PLACE). Multiple alignment of GDPD orthologs in rice was performed via Clustal Omega. Secondary motifs (α-helices, β-strands and 3_10_ helices) were annotated with EndScript (https://espript.ibcp.fr/ESPript/ENDscript). The amino acid sequence of OsGDPD13 was searched against Plant-mSubP, WoLFPSORT, and TMSOC web servers to pinpoint potential signal peptides and transmembrane domains. Possible GPI-anchoring residues in the C-terminal were predicted via the NetGPI (https://services.healthtech.dtu.dk/service.php?NetGPI-1.1) server. We also used MIB2 (http://bioinfo.cmu.edu.tw/MIB2/) to detect putative divalent cation (Ca^2+^, Mg^2+^, Mn^2+^ or Co^2+^) coordination sites.

Novel tertiary protein structures were created from sequence *via* an implementation of ColabFold on Jupyter Notebook (Mirdita et al., 2022) and directly from the AlphaFold2 (AF2) Database (https://alphafold.ebi.ac.uk/). A prediction was performed by the I-TASSER web server by the Zhang Lab Group (https://seq2fun.dcmb.med.umich.edu//I-TASSER/) for comparison and consensus analysis. A survey of productive binding cavities was done using the 3V server (Voss & Gerstein, 2010). Substrate docking poses and binding energy were simulated with AutoDock Vina v.1.2.3 and AutoDock 4.2.6 (Scripps Research Institute, United States). MGLTools and UCSF Chimera 1.15 were also used to prepare the raw PDB structures for modeling (adding Kollman charges, polar hydrogens, etc.) and to visualize enzyme-substrate interactions. Hydrated docking was carried out with the help of the Python scripts and water forcefield distributed by (Forli et al., 2016). Chemical compounds to be docked were downloaded from PubChem or generated from the structure in ChemOffice.

### Generation of *osgdpd13* knockout mutants

Two CRISPR/Cas9 vectors containing dual gRNAs were constructed. The first one targets exon 1 and 2 and the later targets exon 3 and 4. The gRNAs were designed and selected using webservers: https://crispr.dbcls.jp/ and http://rna.tbi.univie.ac.at/cgi-bin/RNAWebSuite/RNAfold.cgi. The cassette containing sgRNA under the control of the pU6 promoter was synthesized and cloned into the cloning vector (pUC57). The gRNA cassette from the pUC57 vector was sub-cloned into vector pOsCas9 by Gateway LR-Clonase (Invitrogen). The CRISPR/Cas9 vector was transformed into *Agrobacterium tumefaciens* strain AGL1, next transformed to Kitakee following the protocol of the International Center for Tropical Agriculture (CIRAD, France) established for the *Oryza sativa* japonica rice.

Transgenic plants and their progenies were genotyped by a PCR-based method. The fragments containing the target site were amplified from genomic DNA by PCR analysis using specific primers flanking the targeted site OsGDPD13-3F (5’-AGTTGTATTGAGCATCTGGA-3’) and OsGDPD13-1R (5’-GGGAACCATTATACCTGAAGCAA-3’). The amplicons were analyzed for induced mutations by electrophoresis on 1.5% agarose gels. The induced mutations were further screened by heteroduplex analysis on native polyacrylamide gel electrophoresis (PAGE) (Zhu et al., 2014). The generated mutations were further validated and characterized by Sanger sequencing using the Sequencing Kit (Applied Biosystems) on the ABI 3100 machine and analyzed by the MUSCLE 3.8.31 program.

### OsGDPD13 subcellular localization

Total RNA was extracted from rice young spikelet using TRIzol @ reagent (Applied Biosystems) as per the manufacturer’s protocol. cDNA was synthesized with a high-capacity cDNA reverse transcription kit (Applied Biosystems) according to the manufacturer’s instructions. The coding sequence of OsGPDP13 without stop codon was amplified from spikelet cDNA using the primers 5’-AAAAAGCAGGCTTCATGCGCCTCTTTACAGCCAA and 5’-agaaagctgggtgTTACACTAGTACGGAAACTA, cloned to entry vector pDONR201 (Invitrogen, USA) and sequenced. The gene was then shuttled to GFP tagging destination vector pH7FWG2 (Karimi et al., 2002) using Gateway LR Clonase (Invitrogen, USA). The obtained vector was introduced to *Agrobacterium tumefaciens* GV3101 to express transiently in *Nicotiana benthamiana* by infiltrating (Ma et al., 2012). 0,1% aniline blue was infiltrated into the leaf, and the leaf was incubated in the dark for 1 hour before imaging. The GFP and aniline blue signals were observed using a confocal microscope (K1-Fluo, Nanoscope Systems, Daejeon, Korea) under 488 nm and 405 nm excitation and detected with a band-pass filter at 520-550 nm and 475 -550 for GFP and aniline blue, respectively.

### Phenotyping the selected mutant lines

Seeds of Kitaake (*Oryza sativa* L.) and three knockout mutant lines were surface sterilized and germinated on ½ MS medium containing 3.0% (w/v) sucrose. Three days after germination (DAG), seedlings were transferred to ½ MS without P to examine P-deficient response. A ½ MS medium containing 625 µM Pi was used as a control. The plants were cultured in a chamber growing at 28/ 26oC with the 14 h:10 h light: dark cycle, 80% humidity, and 12,000 lux light intensity. At 21 DAG, the crown roots that arise from the stem apart from the primary root were manually counted, and the leaves were collected and immediately frozen in liquid nitrogen to analyze the phospholipid content. The plantlets were further cultured in the sand pots under control and Pi starvation treatment conditions for three weeks to record plant height at 42 DAG.

### Lipids extraction and TLC separation

The phospholipids extraction protocol was adapted from (Wang & Benning, 2011) with minor modifications. 550 mg of 21-day-old rice leaves were ground in liquid nitrogen and transferred into glass tubes with a Teflon cap. The crushed leaves were added 5.5 mL of chloroform/methanol/formic acid (10/20/1, v/v/v), and the mixture was agitated at 1300 rpm (4 °C) to dissolve the lipids, then added 2750 μL of 0.2 M H_3_PO_4_ and 1 M KCl solution. Tubes are then centrifuged at 8000 rpm at room temperature, and the lower lipid-rich chloroform layer is collected into glass vials for TLC loading. TLC plates are pre-derivatized by soaking in 0.15 M ammonium sulfate, dried, and baked at 120 °C in 2,5 hours on the experiment date. The mobile phase used in TLC separation was 90 mL of acetone/toluene/water (64/21/5, v/v/v). Upon finish, separated lipids layers were stained yellow using a small closed iodine chamber. Silica-adsorbed lipid bands were scraped off separately into glass vials with PTPE-lined caps.

### FAME reaction and GC-MS analysis

Pentadecanoic acid (C15:0, 2.5 μg/μL) was added to the vial as an internal standard. Lipids are hydrolyzed and their fatty acid tails are trans-esterified into methyl ester derivatives by incubating with Boron trifluoride (14% v/v in methanol) at 80 °C. After one hour, 2 mL of hexane was added, followed by 2 mL of saturated NaCl solution. The vials are centrifuged at 1500 rpm for 5 minutes, and the upper hexane fraction (containing FAMEs) is transferred into a small vial after forcing through a 0.22 μm hydrophilic nylon filter. The solution was subsequently allowed to evaporate under gentle N_2_ until dryness, then redissolved in 100 μL hexane for GC-MS (ISQ-7000, Thermo Scientific, USA). The GC run sequence starts with an initial hold at 110 °C for 2 minutes, then the temperature was raised to 200 °C (rate of 10 °C/min), 250 °C (4 °C/min), and finally 280 °C (20 °C/min). The percentage of molar of FAMEs and lipids is calculated according to the following equations (Wang & Benning, 2011):

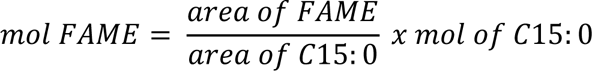

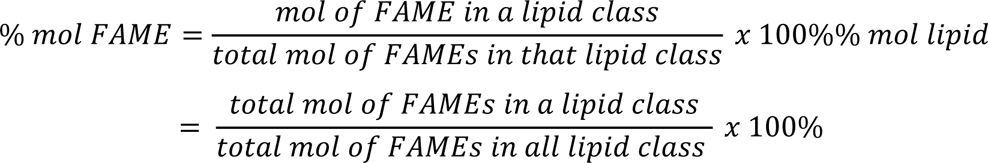

### Statistical analysis

The data represented the mean ± SD from 20 seedlings. The statistical analysis (ANOVA, *t*-test) and graphing were conducted using GraphPad 10.0.

## RESULTS

### *OsGDPD13* is absent from chromosome 9 of the *Indica* rice’s genome

The raw sequencing data from the *Japonica* cultivar (G299) showed a considerably greater number of matched reads to the *Japonica* OsGDPD13 promoter sequence, which covered nearly 100% of the promoter region. In contrast, the three *Indica* varieties (G22, G99, and G207) exhibited gaps from 100-650 bp, 750-950 bp and no detected reads from 1350^th^ bp downwards **(Fig. 1A)**. Detected reads in *Indica* samples demonstrated discernable differences from that of the reference sequence.

**Figure 1.**
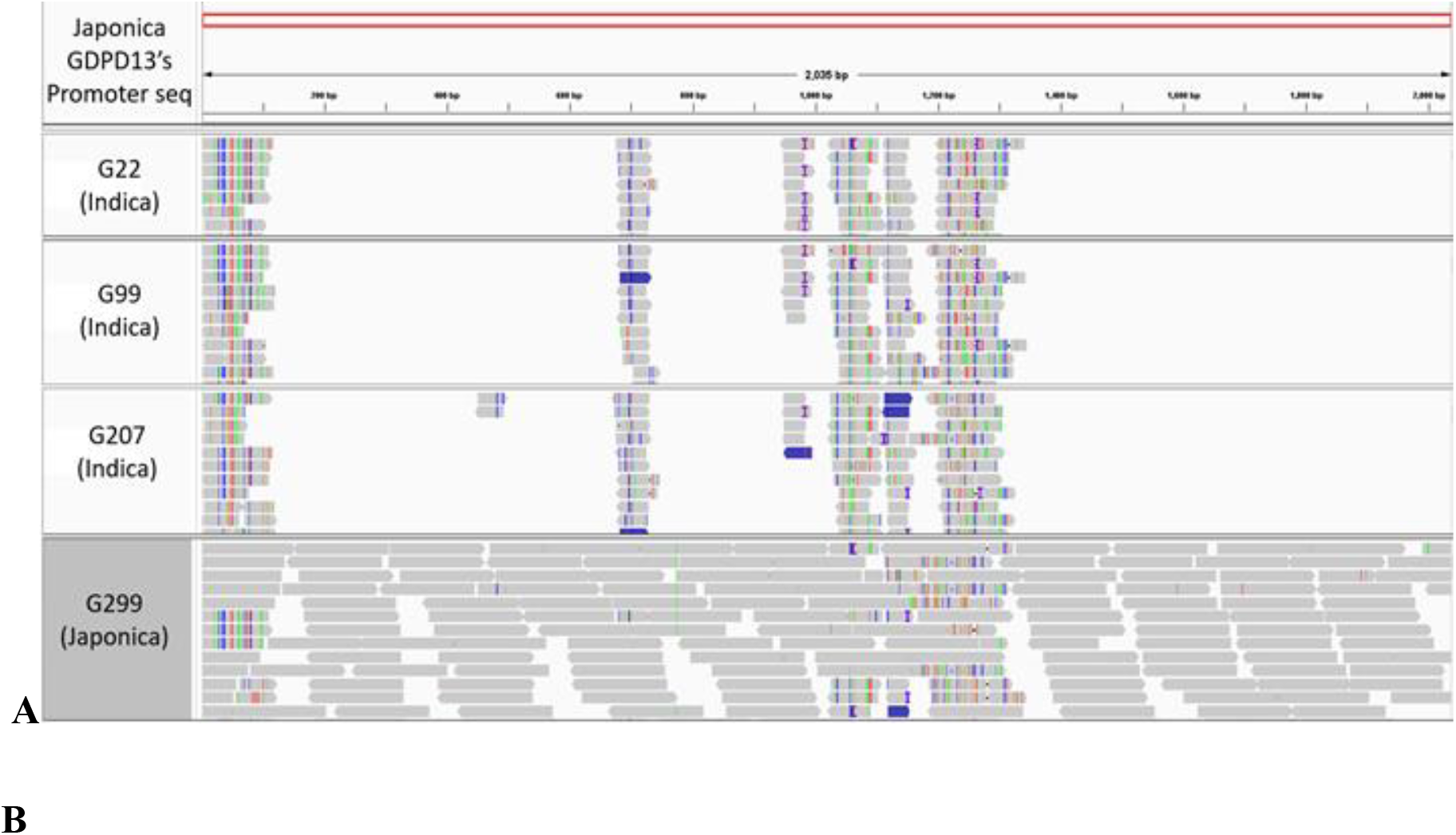
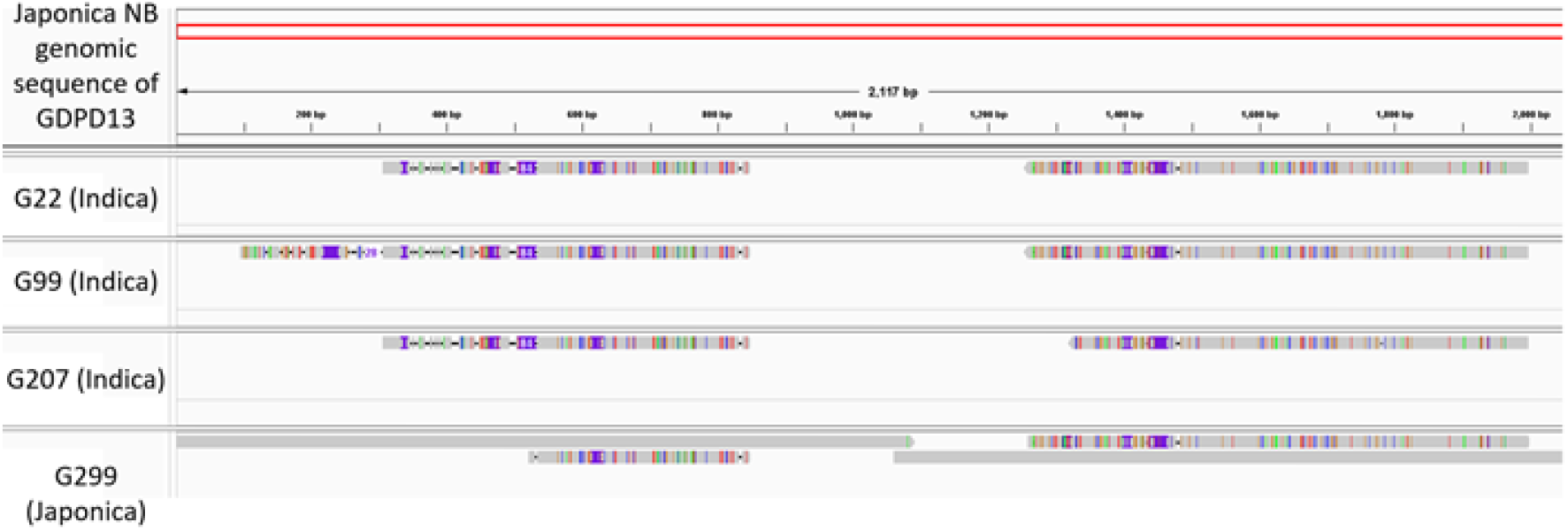
Alignment of *Indica* (G22, G99, G207) and *Japonica* (G299) to OsGDPD13 *Japonica* reference. (**A**) Raw reads were mapped to the OsGDPD13 promoter region. (**B**) Contigs were mapped to the GDPD13 in the *Japonica* reference genomic region. Visualization of these alignments was performed on IGV. Red bars denote reference sequences with the scale below. Grey fragments exhibit matched Illumina short reads/contigs to the reference regions. Multiple colors within these reads/contigs indicate nucleotide changes compared to the reference sequences.

Next, the Illumina short reads from the four rice samples mapped to the OsGDPD13 *Japonica* reference genomic sequence were assembled into contigs. The *Japonica* G299 had four contigs matched to the OsGDPD13 reference gene, two of which perfectly matched the reference, and two contigs showed partial matches with significant variability in nucleotide positions. On the other hand, the other three *Indica* samples (G22, G99, G207) had only two contigs, which partially covered the full-length OsGDPD13 gene. These contigs also exhibited significant divergence from the *Japonica* reference sequence as there existed many color bands indicating nucleotide changes. Noticeably, the contigs from the three *Indica* samples also showed gapped band patterns similar to the two dissimilarity contigs in the *Japonica* sample (G299) **(Fig. 1B)**. To further investigate the origin of these contigs, we performed BLASTn homology search against NCBI’s non-redundant (nr) database, and we identified that the three homologs to each contig were located on chromosome 12 of *Indica* and *Japonica* rice with > 99% identity and query coverage of 100% and encode for OsGDPD7, the closest gene to OsGDPD13 in OsGDPD family. In addition, alignment of the OsGDPD13 gene with available *Indica* genomes in the NCBI GenBank database showed that OsGDPD13 discontinuous matched to several regions from the identical subject sequences **(Fig. S1A)** and an area at chromosome 12 of the *Oryza sativa indica* cultivar RP Bio-226 displayed the highest similarity with 82.66% identity and 52% query coverage **(Fig. S1B)**. Altogether, the *in-silico* investigation results showed *OsGDPD13’s* absence from the genome of *Oryza sativa Indica* rice plants, as the examination was done on both *OsGDPD13’s* promoter and genomic sequences.

### The tertiary structure of OsGDPD13 showcases motifs common to GDPDs

For the OsGDPD13 protein, ColabFold generated a total of twelve β-strands, twelve α-helices, and six 3_10_ helices, three of which fall in the GDPD-insert (GDPD-I) domain, a unique structure for the GDPD subfamily. The core structure consists of eight strands (β1, β2, β5, β6, β7, β10, β11, β12) in alternation with eight helices (α2, α3, α5, α6, α8, α10, α11 and α12) (Fig. S5) forming a (α/β)_8_ TIM-barrel tertiary fold, concurring with the structures of other OsGDPDs. The presumed GDPD-I fold is located in the loop between β2 and α3. The superimposition of the two OsGDPD13 MSU structures shows the great fit of the TIM-barrel between independent predictions of I-TASSER and ColabFold **(Fig. 2A, B)**. As the literature suggests that the active site of GDPDs is near the center of the core with only a few residues on flexible loops (Myers et al., 2016), we opted to use this prediction by ColabFold as a framework for further docking analyses. Remarkably, the pIDDT (a local score for comparing distance) given for MSU structure by ColabFold demonstrates high confidence in the core region except for the previous region comprising α1, α2, and β1.

**Figure 2.**
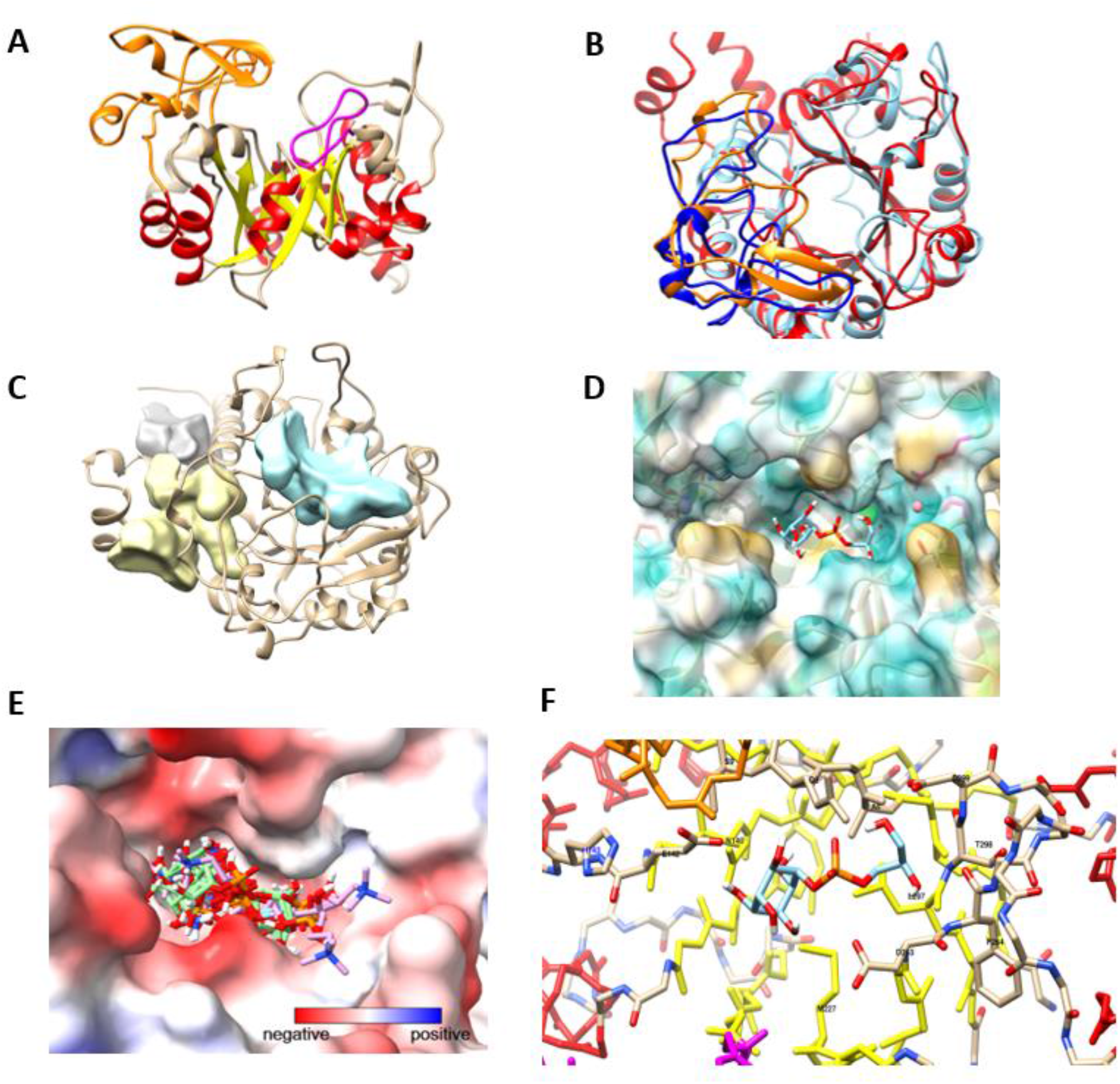
(**A**) OsGDPD13 protein by ColabFold and **(B)** Superimposition of ColabFold (red) and I-TASSER (blue) tertiary structure (**C**) Possible cavities of OsGDPD13 (**D**) Hydrophobic surface generation of docked GPI to OsGDPD13, along with the projected positions of four metal ions (Ca^2+^ (green sphere), Mg^2+^ (yellow, hidden inside transparent surface, left), Mn^2+^ (purple hidden, left) or Co^2+^ (red, right) (**E**) Electrostatic potential mapping of OsGDPD13, with clustering of all binding poses by GPC, GPE and GPI revealed (**F**) Backbone trace of OsGDPD13, assumed important residues interacting with GPI denoted.

The TMSOC (Transmembrane helix: Simple or Complex) web server automatically detected a transmembrane domain (twilight-TM) between residues 38 and 60 (hydrophobicity score of 3.6). This is somewhat supported by the low hydropathy index and solvent accessibility stretch from amino acid 51 to 59. This result coincided with the Interpro analysis that GDPD13 contains a transmembrane domain at the N-terminus and just sited before the PI-PG domain responsible for the phosphoinositol phosphodiesterase’s activity. Nevertheless, another transmembrane region was manually defined from 301 to 325 comprising the α12 helix to the C-terminal, which the server returns as a complex-TM (hydrophobicity score -1.82). Net-GPI 1.1 found no GPI-anchoring residues (ω-sites) in the C-terminal (chance of site 26.7%).

### OsGDPD13 might have an affinity for GDPD substrates

As the literature suggests that the active site of GDPDs is near the center of the core with only a few residues on flexible loops (Myers et al., 2016), we opted to use this prediction by ColabFold as a framework for further docking analyses. The binding pocket of OsGDPD13 (in cyan) **(Fig. 2C)** possessed a volume of 1209 Å^3^ as estimated by the 3V server, which is 10 times larger than that of bGlp from *B. subtillis*. Nevertheless, the real exposed region for binding is possibly lesser as the position of the flexible chains from a predicted structure cannot be verified. The bGlpQ flexible loop is derived from crystallographic data and, therefore, is expected to be closer to the productive conformation of the substrate. Binding poses of hypothetical substrates **(Fig. 2E)** demonstrate a high degree of clustering at the active cavity, even with a greater search volume. A single histidine residue (His143) in the vicinity of the active site is labeled in blue **(Fig. 2F)**; however, it is not expected to participate in ligand binding due to its great distance. No accessible metal ion binding motif can be found for Mg^2+^ or Mn^2+^ *via* the MIB2 server that does not involve heavy clashing with the binding hole or being completely out of position **(Fig. 2D)**. A Co^2+^ can be nested in the active site but is too distant for catalytic purposes. Only Ca^2+^ (green sphere) appears decently positioned in the cavity.

Reasonable K_M_ values for GDPDs ranged from 0.1-10 mM **(Table 1)**. Hydrated docking has been cautioned to give unreliable binding energy as post-processing is required (Forli & Olson, 2012). Still, when combined with docking results for OsGDPD2 **(Table S1)**, another newly identified GDPD with glycerophosphodiesterase activity, it is possible that taking water interaction into account might at least convey the ranking of substrate selectivity. Projected order of binding affinity for OsGDPD13: GPI > GPC > GPE.

**Table 1.** Binding energy and predicted binding affinity by AutoDock4 and hydrated docking results on OsGDPD13

| Substrate | $\Delta G$ by AD4 | Predicted $K_M$ (mM) | $\Delta G$ by AD4 with water | Predicted $K_M$ ($\mu M$) |
| --- | --- | --- | --- | --- |
| GPE | -2.92 | 7.23 | -4.42 | 580.03 |
| GPC | -3.02 | 2.63 | -6.09 | 34.41 |
| GPI | -4.18 | 0.862 | -8.51 | 0.574 |

### GDPD13 protein localizes to the plasma membrane, cytoplasmic speckles, and plasmodesmata

GPDP has been reported to function in different cellular compartments. Hence, the subcellular location prediction of OsGDPD13 was first investigated by Plant-mSubP. Using pSORTb sub-location prediction results showed that OsGDPD13 might target the cytoplasmic membrane. Therefore, the subcellular localization was validated using a GFP tag fused at the C-terminal of the protein under 35S promoter. In this study, the coding sequence of OsGDPD13 was amplified from a cDNA library made from mRNA isolated from young flowers of *Oryza sativa* ssp—japonica *variety* Dongjin. GDPD13 ORF consists of 1167nt, coding a protein of 389aa. GFP signals were recorded at the early expressing stage at two days after infiltration to the late expressing stage at four days after infiltration. The results showed different localizations of OsGPDP13, including plasma membrane (Fig 3A), plasmodesmata (Fig. 3B), and cytoplasmic speckles (Fig. 3A). Closer view of the signal on the plasma membrane denotes punctate and patched fluorescence which resemble plasmodesmata (PD). Therefore, aniline blue, which binds to callose (beta 1,2 glucan polymer), was used to confirm OsGDPD13 localization in PD (Fig. 3B). This diverse location was further confirmed by the retained signals on the cell wall observed after plasmolysis (**Fig. 3C**).

**Figure 3.**
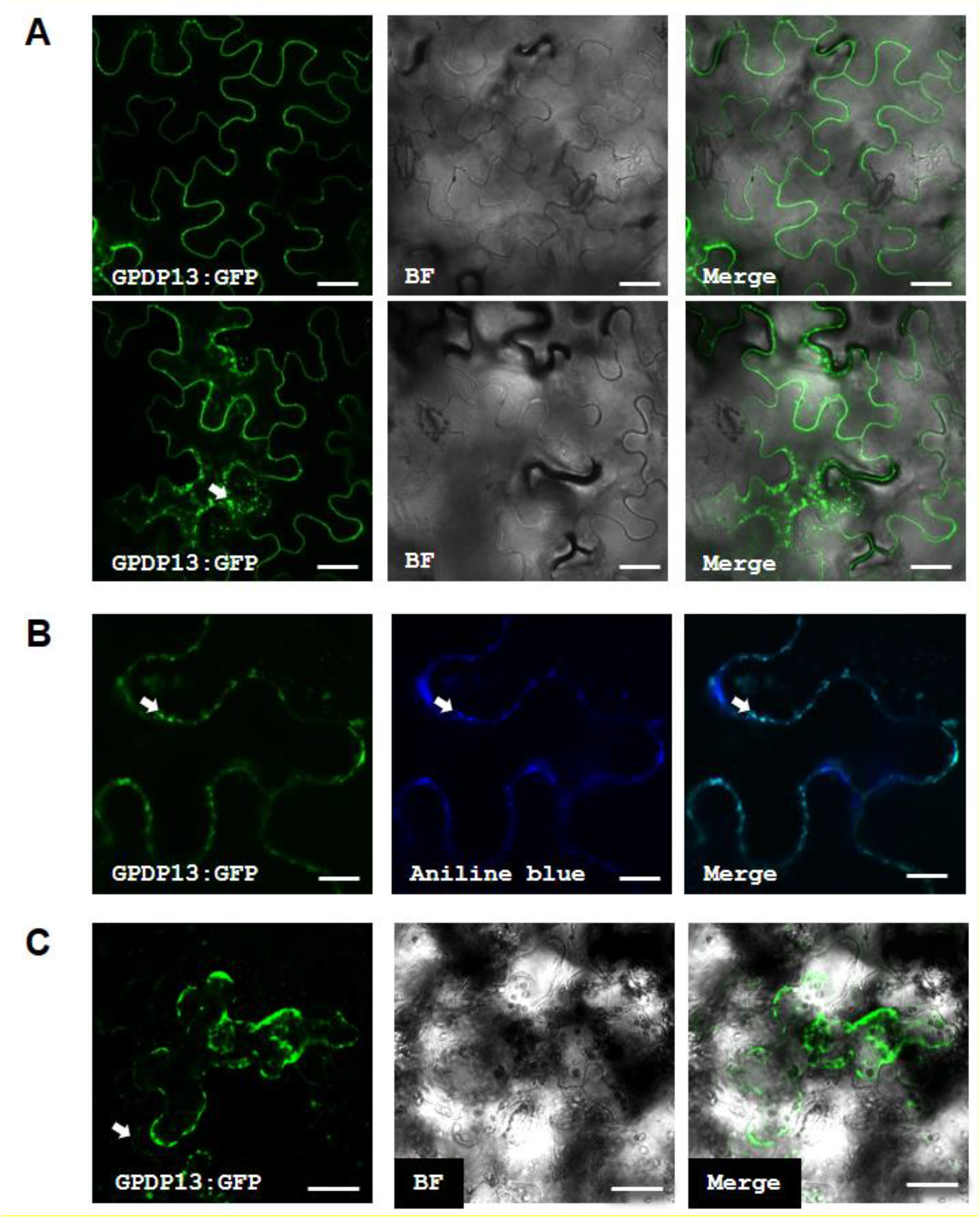
Image of GPDP13::GFP in *Nicotiana benthamiana* leaves four days after infiltration by confocal laser scanning. (**A**) GFP signals were detected in the plasma membrane (upper row) and cytoplasmic speckles (middle row, red arrow). (**B**). Callose deposition in plasmodesmata was validated by aniline blue staining. Arrows indicate co–localization of OsGDPD13 and aniline blue signals. (**C**) GFP signals 15 mins after plasmolysis by NaCl 1M. An arrow indicates retained GFP signals on the cell wall. Bar = 30 µm (A and C), 10µm (B). BF: bright field.

### Induced mutations of OsGDPD13 gene

The osgdpd13 mutant lines were generated on the Kitaake variety belonging to the O. sativa Japonica genetic background for loss of function study using CRISPR/Cas9 **(Fig. 4)**.

**Figure 4.**
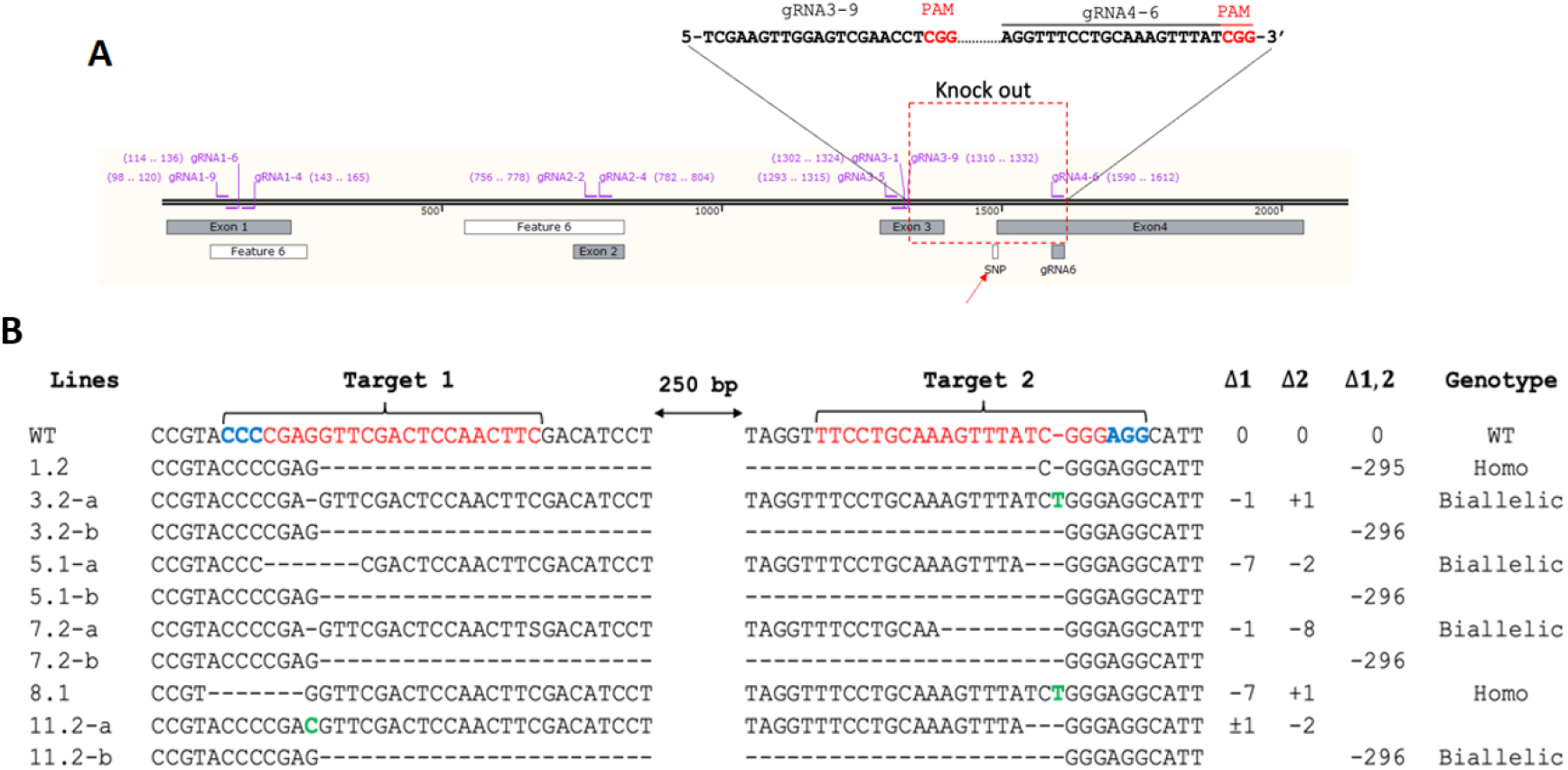
Identification and characterization of induced mutations of the *OsGDPD13* gene at the T0 generation. (**A**). *OsGDPD13* gene structure and locations of gRNA target sequences. **(B)** Sequencing results of the OsGDPD13 mutations in T0 transgenic rice lines. 1.2-11.2: transgenic rice lines. a/b indicates different alleles for each T0 line, red letters indicate sgRNA sequences, blue letters display PAM sites, and green letters show the inserted sequences; Homo and Biallelic are for homozygous and biallelic genotypes, respectively.

DNA extracted from transgenic lines was confirmed deletion mutation by electrophoresis, and expected small indels in the *OsGDPD13* gene were detected using the PAGE-based genotyping assay. PCR amplicons from the six expected mutations of the *OsGDPD13* lines were sequenced by the Sanger method, which further confirmed that all the potential deletion mutations screened in the electrophoresis were specific to the targeted site in biallelic or homozygous states **(Fig. 4B)**. In the T1 generation, the progenies of three representatively edited lines 1.2, 7.2 and 8.1 were assessed in terms of the inheritance and segregation of *OsGDPD13* targeted mutations. Based on the PCR gel electrophoresis and Sanger sequencing results **(Fig. S2)**, we successfully identified potential homozygous mutant lines for the phenotyping analysis at T3 generations.

### Loss-of-function of OsGDPD13 leads to inhibition of growth under Phosphate starvation conditions

To identify the effects of the *OsGDPD13* mutations on the growth of rice plants, three mutant lines (*osgdpd13*.8.1, *osgdpd13.*7.2, and *osgdpd13*.1.2) carrying homozygous knockout alleles were grown for three weeks in the test tube for 21 days and then transferred to sand pots and grown until 42 days under control and Pi-starvation conditions, with Kitaake used as the wild type. At three weeks, the KO mutant seedlings did not inhibit significant differences in shoot length compared with the WT plant under low Pi conditions. Regarding the number of crown roots, the performance of three mutant lines under Pi deficiency conditions was significantly lower within 21 days compared with WT plants (Fig 5A-B). The observable reduction of crown root formation could be considered an abnormal phenotype caused by mutations in the *OsGDPD13*.

**Figure 5.**
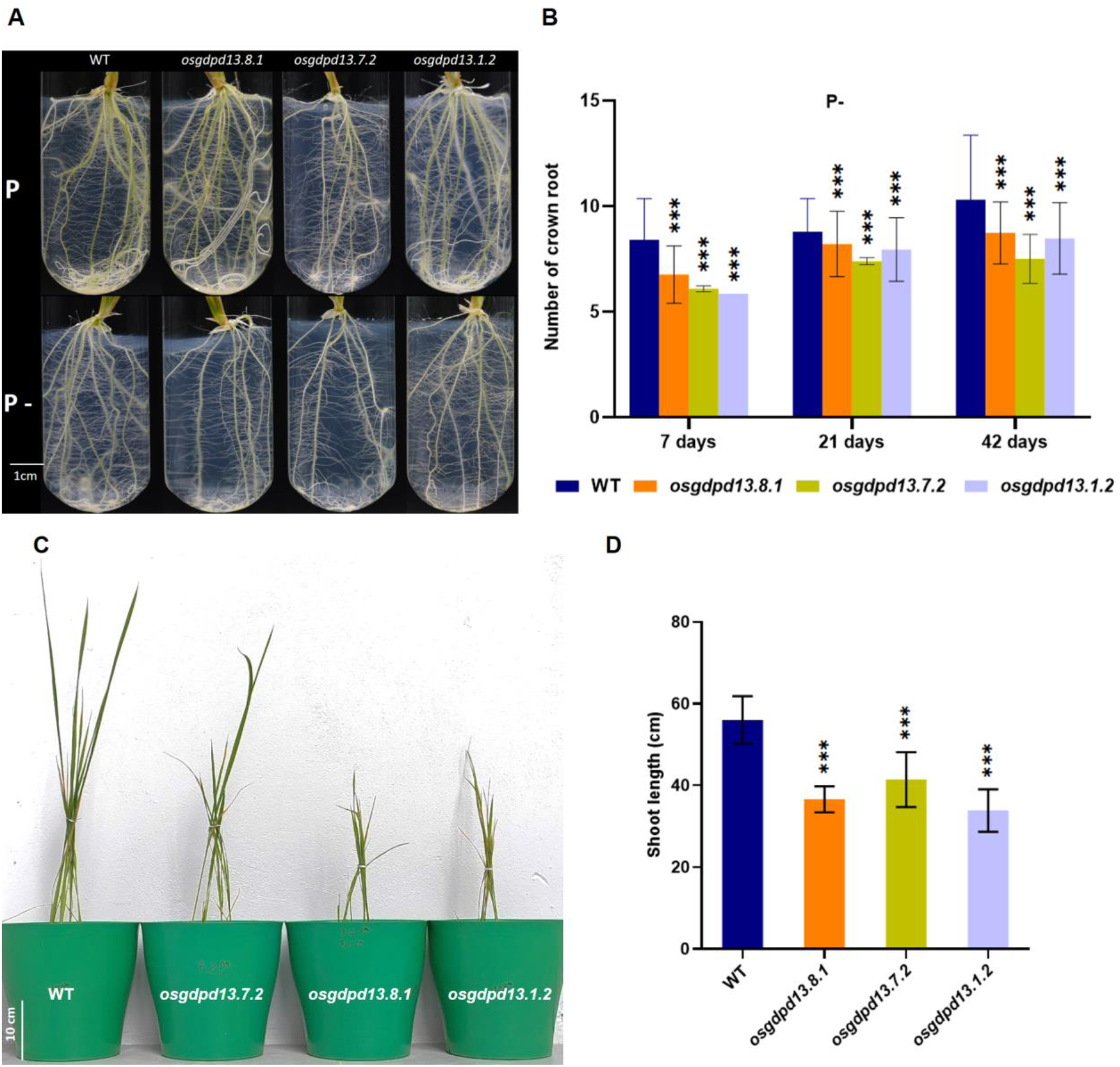
Effects of knocked out *osgdpd13* on the number of crown roots and the shoot length of rice under Pi-starvation condition. (**A**) Crown root phenotype of three *osgdpd13* mutants and WT after 21 days under normal (upper) and Pi-starvation conditions (lower), scale bar = 1 cm. (**B)** The number of crown roots of three *osgdpd13* mutant lines compared to wild-type (Kitaake) after 7, 14 and 21 days under P-starvation conditions. (**C**) *osgdpd13* mutant plants and shoot length **(D)** after 42 DAG under Pi-starvation conditions, scale bar = 10 cm. Data are expressed as mean ± S.D. (n = 20). ANOVA and then Student t-tests were used to analyze the differences between mutant and WT. Asterisks represent different levels of significance (*P<0.05, ***P<0.001).

We further assessed the effects of *gdpd13* KO mutation on the shoot growth of three mutant lines and WT after 42 days grown in low Pi media. All three mutants showed apparent retardation of the shoot growth compared to the WT in response to Pi starvation stress (Fig 4C-D). In general, the *osgdpd13* loss-of-function line exhibited a reduction in shoot length, and this decrease might be due to altered Root system Architecture (RSA) in the mutants during Pi starvation treatment.

### Loss-of-function of OsGDPD13 reduces phospholipid modification in rice leaves under Phosphate starvation conditions

The 21-old-day rice leaves of the wild-type plants and the three *osgdpd13* lines were investigated in phospholipid composition (phosphatidylcholine (PC), phosphatidylethanolamine (PE), phosphatidylinositol (PI), (phospholipid phosphatidylglycerol (PG)); glycolipids (monogalactosyldiacylglycerol (MGDG) and digalactosyldiacylglycerol (DGDG)) and sulfolipids (sulfoquinovosyl diacylglycerols (SQDG)) level under both control and Pi-deficient conditions. In general, glycolipids (MGDG, DGDG) and sulfolipid (SQDG) accounted for the majority of lipid class in both P-deficient and P - sufficient media in the wild-type and the mutant lines (**Fig. 6A-B**). Under control conditions, the phospholipids (PL) content in the wild type was significantly lower than those of the mutant lines, which also means the glycolipids (GL) content was significantly higher compared to the mutant (**Fig. 6A-B**). It is noticeable that the WT line showed a more marked decline in the relative level of total phospholipids compared with the mutant lines. Specifically, a considerable reduction in the content of PL and an increase of GL content was observed in the WT line (80%). In contrast, only 15-25% of PL was recorded in three KO mutant lines*, osgdpd13.8.1, osgdpd13.7.2, and osgdpd13.1.2* mutant lines (**Fig. 6C**).

**Figure 6.**
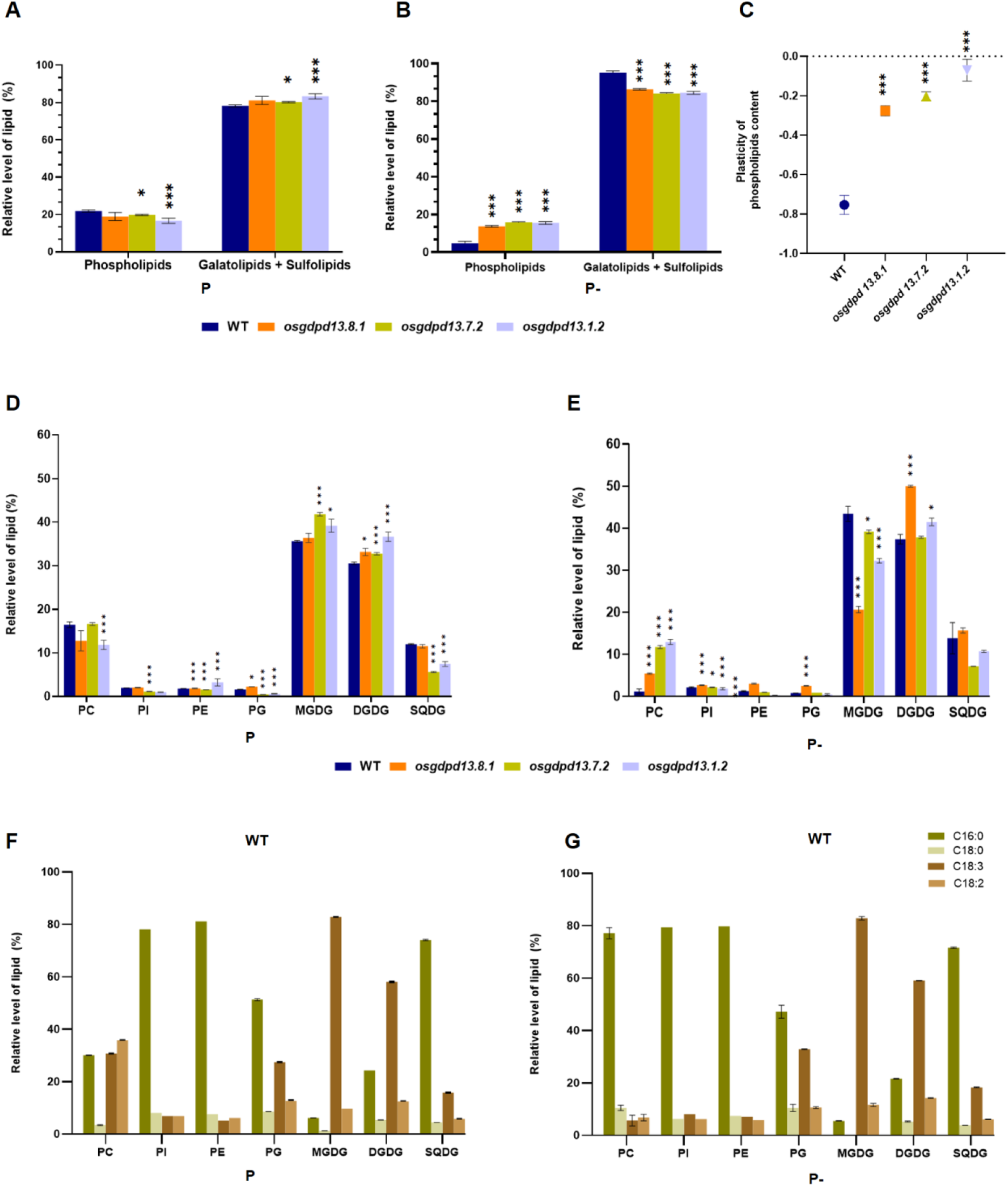
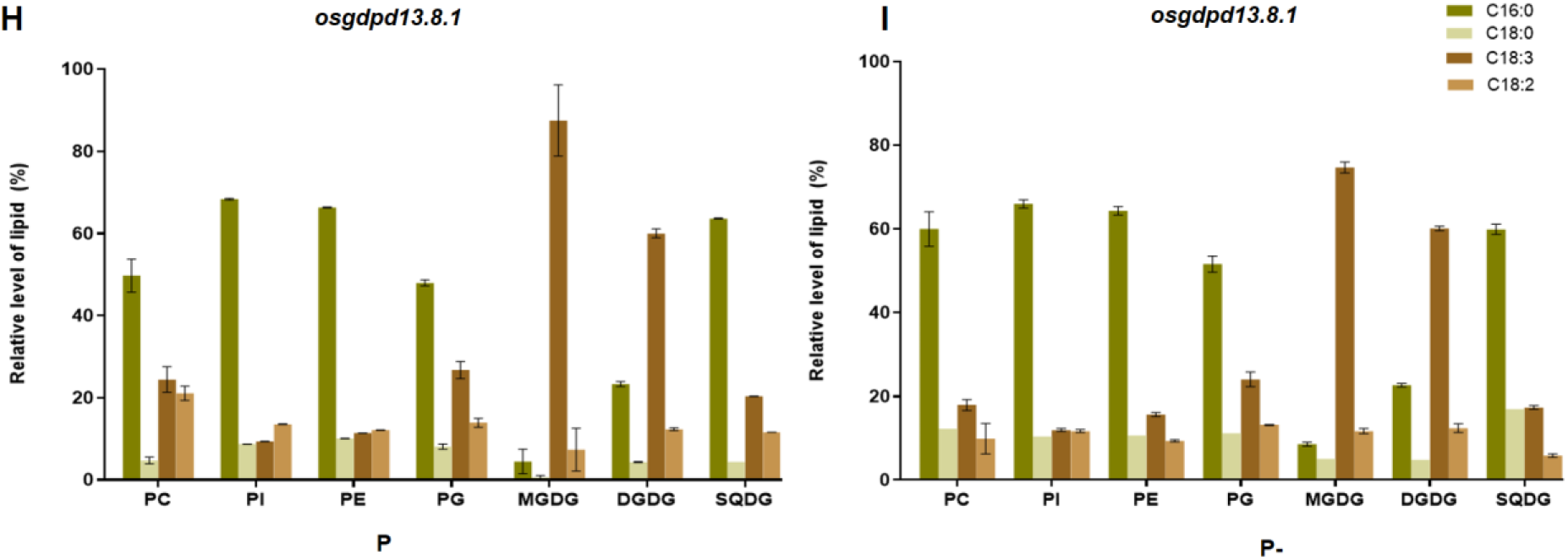
Effect of *OsGDPD13* knocked out on rice leaves lipid profiles after 21 days grown under Pi (P-sufficient) and Pi- (P-deficient) conditions (**A, B**) The total relative content of phospholipid, galatolipid, and sulfolipid of green leaves of WT (Kitaake) and *osgdpd13* mutant **(C)** Plasticity index of phospholipids relative content of WT (Kitaake) and *osgdpd13* mutant lines. (**D, E**) The relative level of lipid class species of WT and osgdpd13 mutant lines (F, G, H, I) Molecular species of a phospholipid, galactolipid, and sulfolipid of WT (Kitaake) and *osgdpd13* 8.1 mutant lines. % mol of lipids was calculated from GC areas of FAMEs. Data are expressed as mean ± S.D.; asterisks represent different levels of significance (*P<0.05). Data are expressed as mean ± S.D. (n = 3).

Regarding lipid species, PC and DGDG content were remarkably higher, while the MGDG content was much lower in the mutant lines than in the WT in low Pi conditions. Pi starvation leads to the reduction of MGDG while increasing DGDG in mutant lines compared to WT rice. Besides, the decline of PC and PG content and the increase in DGDG content were the most apparent differences between WT and mutant lines. While a significant decrease in PC and PG levels was found in the WT, these corresponding levels of mutants slightly fell in response to Pi deficiency (**Fig. 6D, 6E, Table S2**).

Regarding molecular species of each lipid class, there was no significant difference in fatty acid species and relative proportions between species among WT lines and mutant lines grown in both Pi-supplied conditions after 21 days. Fig 6F and 6G showed that under normal and Pi starvation stress, leaves rice extracted from WT and mutant lines consisting of 4 main fatty acid species, including Palmitic acid (C16:0), Linoleic acid (C18:2), Linolenic acid (C18:3) and Stearic acid (C18:0). In all lines, under the control and treatment conditions, C16:0 made up the main part of phospholipids and sulfolipid class, the three remaining fatty acids only accounted for a small proportion (**Figure 6E-I**). Meanwhile, C18:3 is the primary glycolipid component in both WT and mutant lines under two conditions.

To summarize, there was a decrease in total phospholipid content and a significant rise in both glycolipid and sulfolipid relative content in both the WT and mutant lines to deal with Pi starvation stress. It is worth considering that the partial replacement of phospholipids by glycolipids and sulfolipids of the wild-type line is significantly stronger than that of the three *osgdpd13* loss-of-function mutant lines in response to Pi-deficient conditions.

## DISCUSSION

### OsGDPD13 shares a common tertiary structure motif to the GDPDs family and might have a weak affinity for GDPD substrates

OsGPPD13 encodes GLYCEROPHOSPHODIESTER PHOSPHODIESTERASE (GDPD) enzyme, which is involved in the degradation of glycerophosphodiesters into G-3-P - an important metabolite in maintaining phosphate homeostasis (Ohshima et al., 2008). Most members of the OsGDPD family (from *OsGDPD1* to *OsGDPD12*) are well-established under different nutrient stresses (N, P, Fe, K), with *OsGDPD2*, in particular, being correlated with phospholipid remodeling and synthesis (Mehra & Giri, 2016). However, *OsGDPD13* remains under-explored due to a lack of cDNA database in several reports. There is little information regarding its expression except in microarray transcriptomic data suggesting mild upregulation in the root and panicle of rice under P stress. This is compounded by conflicting intron, exon, and start site annotations in independent rice databases such as MSU and RAPDB. The protein evidence record for OsGDPD13 (NP_001062919) was removed from the RAPDB database in 2021 due to standard genome annotation processing. To and colleagues (2020) observed among 157 Vietnamese rice landraces, the *O. sativa* subsp. *indica* subgroup was strongly affected by low Pi treatment than japonica, as demonstrated by three significantly different relative roots to shoot ratios, the relative efficiency of P use, and relative physiological PUE traits (To et al., 2020). The results suggested that phenotypic heterogeneity among *indica* and *japonica* subpopulations under Pi starvation conditions comes from diversity in genetic backgrounds. Interestingly, our result indicated that a genomic region was missed on Indica (G22, G299, G207) genomes as well as OsGDPD13 is absent from the genome of *Oryza sativa Indica* rice subgroup when their draft genomes aligned back to Japonica OsGDPD13. This may suggest that OsGDPD13 contributes to adapting *japonica* to phosphate deficiency.

*In silico* analysis of OsGDPD13 demonstrates relatively high structural similarity with other enzymes in the “functional” GDPD family, as I-TASSER was able to recognize *E. coli* GlpQ by TM-align (TM-score 0.865) as a template for constructing its 3D structure despite low sequence identity (0.170) because no crystal structure for plant GDPDs (whether *Arabidopsis* or rice) is presently available. However, the TIM-barrel core, in general, is appropriated by many enzyme families with admirable diversity and is thought to show highly divergent evolution (Kadumuri & Vadrevu, 2018). Researchers have associated this fold with substrate binding capability, favoring a phosphate chelating function probably as a consequence of the original (triose phosphate isomerase) enzyme fold (Wierenga, 2001).

In fact, the top prediction for OsGDPD13 function by COACH was a sucrose glucosylmutase (EC 5.4.99.11), although the following consensus confirmed the GDPD classification (EC 3.1.4.46). Gene Ontology (GO) terms in the I-TASSER database and UniProt automatic annotation did identify the eponymous GD-PDE domain (itself a subclass of the parent PI-PLC - phosphoinositide phospholipase C family). OsGDPD13 featured a single complete GD-PDE domain, also predicted internally to have no signal peptide and a “twilight” TM region **(Fig. S5),** which means OsGDPD13 falls between a complex-TM (likely to be involved in function) and simple-TM (a helical anchor). Upon closer inspection, OsGDPD13 does not possess any motif of the two related phosphatase families (purple acid phosphatase or GDPD), which are the (PD-PNH) (Koonin, 1994) and the GDPD signatures (HR(X)_n_EN(X)_n_EXD(X)_n_HD) (Jackson et al., 2007), where X stands for any amino acid. In contrast, OsGDPD2, a newly functional enzyme of the GDPD family, recaptured the latter motif with fidelity (Mehra et al., 2019). The PI-PLC family in general and identified functional GDPDs from bacteria and plants in particular consists of two His residues in the catalytic cavity acting as proton acceptors and proton donors in catalyzing the hydrolysis of the phosphodiester bond (Yip et al., 2011) OsGDPD13 possesses a single histidine residue (His143, Fig. 5D) close to the active site discovered *via* docking. Phylogenetic analysis has segregated OsGDPD13 into a clade along with AtGDPDL1 and SHV3 distinct from active GDPD enzymes with conserved active sites, which clustered into two families (cytosolic UgpQ and periplasmic GlpQ), with corresponding alignment showing little sequence identity (Mehra & Giri, 2016). While SHV3 did not display hydrolysis function toward GPC, AtGDPDL1 had a weak affinity to GPC, GPE, and GPI using Ca^2+^ as a cofactor (Cheng et al. 2011b; Hayashi et al. 2008). Surprisingly, like OsGDPD13, AtGDPDL1 also had only His184 adjacent to the active site **(Fig. S4)**. Thus, while the catalytic mechanism theoretically featured two histidine residues **(Fig. S3),** a single residue may be sufficient to hydrolyze the phosphodiester bond, as reported in the case of AtGDPDL1 (Cheng et al. 2011b). For metal binding loop prediction, OsGDPD13 has no accessible binding motif for Mg^2+,^ Mn^2+^ but Ca^2+.^ Serving as a metal cofactor, Mg ^2+^ promotes plant GDPD activity more than Ca ^2+^, although Ca ^2+^ probably assisted catalyzing to a minor degree (Cheng et al., 2011b; Mehra et al., 2019; Mehra & Giri, 2016). Interestingly, Mehra et al., 2016 reported that CaGDPD1 had activity even without divalent cations. Previous studies reported bacterial GDPD displayed the difference in divalent cofactor preference. At the same time, glpQ activity was restored by Ca2+ or Cd2+ Mn2+, Cu2+, Co2+, Mg2+, Zn2 in the order of priority (Larson & van Loo-Bhattacharya, 1988), UgpQ required Mg2+, Co2+, or Mn2+ for its enzyme activity (Ohshima et al., 2008). Regarding substrate affinity, docking evidence suggested that OsGDPD13 has an affinity for GPI. By comparing Km and delta G values with functional enzyme OsGDPD2 and GlpQ, UlpQ in the same docking model, GPI can be considered a strong substrate for putative active GDPD13. Taken together, although sharing a homologous physical structure to the GDPDs family and being predicted to have an affinity to GPI, OsGDPD13 does not possess typically catalytic features such as a conserved active site and a – metal-binding loop (except for Ca2+) compared to 2 bacterial functional GDPD (glpQ, ugpQ) and plant functional OsGDPD2. This suggests that OsGDPD13 might have weak catalytic activity.

### Loss-of-function of OsGDPD13 reduces root growth and leaves phospholipids content in response to Phosphate starvation condition

It has been observed that P deficiency significantly induces GDPD expressions in many plants, and the knockout mutants displayed a similar pattern of slowed root development under the P deprivation (Cheng et al. 2011b; Mehra et al. 2019; Mehra and Giri 2016). In this work, the functional characterization of the gene *OsGDPD13* in response to low Pi was performed in a *Japonica* rice variety, Kitaake. The differential root morphologies between WT and *osgdpd13* mutant plants were observed after 21 days of treatment. However, the significant difference in shoot length between WT and 03 knockout mutant lines was merely detected after 42 days under Pi starvation. Specifically, plant height was observed to decrease in the KO *osgdpd13* mutation plants. This short shoot phenotype may just be a consequence of the alteration of root system architecture under Pi-deficiency stress. Besides, phospholipids were partially replaced by glycolipid and sulfolipid in both WT and *osgdpd13* KO mutant; especially, the proportion of phospholipid modification (especially PC, PG) of loss-of-function lines was significantly lower than that of WT. The *osgdpd13* knockout mutants were similar in terms of retarded root growth under P starvation, proposing the important role of this candidate gene in modifying the root system architecture. These results also suggested the involvement of OsGDPD13 in phospholipids modification during membrane lipid remodeling.

Compared to the identified *OsGDPD* genes, a number of GDPDs have been reported to be involved in root development or in Pi recycling to adapt Pi deficient stress. Most type A *OsGDPDs* except *OsGDPD4* showed early response with significantly upregulated expression to low P treatment (7 days). In comparison, most of the type B *OsGDPDs* (*OsGDPD7* to *OsGDPD13*) were late responsive (after 15 days), preferentially in roots (Mehra et al., 2019). Regarding root and shoot performance, only three *OsGDPDs* (*OsGDPD2*, *OsGDPD3*, and *OsGDPD5*) were induced in shoot tissue under Pi deprivation (Mehra & Giri, 2016). According to Mehra et al., 2019, *OsGDPD2* played a crucial role of *OsGDPD2* in root growth regulation under low Pi levels. OsGDPD2 affected root biomass and root length but not lateral root length and density. In *Arabidopsis*, *AtGDPD6* plays a role in root length, and lateral root formation did not show any significant differences in shoot morphology compared to WT under Pi deficiency conditions (Ngo & Nakamura, 2022). Regarding lipid membrane remodeling, OsGDPD2 induced both galatolipids and phospholipid biosynthesis under Pi starvation and might play a potential role of OsGDPD2 in *de novo* glycerolipid biosynthesis or Kennedy pathway (Mehra et al., 2019), two another GDPDs, the AtGDPD1 and AtGDPD6 did not affect membrane lipid composition in Pi-starved conditions (Cheng et al. 2011b; Ngo and Nakamura 2022)).

Up to present, there is still a gap of insight explanation in the molecular mechanism by which plant GDPDs family regulate in response to Pi starvation stress. Phospholipids degradation can be catalyzed by two pathways involving non-specific PLC (NPC4/5) or PLD (S. Zhu et al., 2022). The third pathway by which glycerophosphodiesters (derived from the deacylation of phospholipids) are hydrolyzed to form G-3-P by GDPDs has not been clarified(S. Zhu et al., 2022). Previous studies proposed that root growth was influenced by downstream products of GPDG hydrolysis, such as G-3-P and PA. GPDD hydrolyses GDP to form G-3-P, which can serve as a precursor of PA. PA could be incorporated into DAG, which could be used for lipid synthesis or further degraded into inorganic phosphate (Pi). PA acts as a signal molecular in response to different nutrient deficiencies (Wang et al., 2006). Cheng et al. 2011b reported that G-3-P could rescue the decreased root growth phenotype of the KO *atgdpd1* under Pi starvation stress. Similarly, impaired root growth of *Atgdpd6* mutants under phosphate starvation was recovered by adding glycerol 3-phosphate but not with choline (Ngo & Nakamura, 2022)). Both *Atgdpd1* and *Atgdpd6* have enzymatic activity and were proposed to be not involved in *de novo* phospholipid biosynthesis pathway due to unaltered lipid composition under low Pi stress, but instead involved in G3P homeostasis in root growth (Cheng et al. 2011b; Ngo and Nakamura 2022)). Meanwhile, (Mehra et al., 2019) proposed a potential role of a functional OsGDPD2 in *de novo* lipids biosynthesis or Kennedy pathway by which the accumulated PA enhanced root growth.

However, our results showed that the silencing of OsGDPD13 caused a significant change in root formation and in modification of phospholipids level despite the fact it may be inactive or have very weak activity through *in silico* protein prediction. This suggests that this gene is involved in the alteration of phospholipids, whether it has catalytic activity or not. Previously, many studies reported that an inactive enzymatic protein may regulate the catalytic mechanisms of the active ones in its family in various ways. They increase/ decrease the catalytic efficiency by allosteric regulation through (1) the formation of heterodimers or heterooligomers, or (2) molecular switches in signaling pathways, (3) restraining ligands in specific locations, or (4) serving as competitive inhibitors thanks to their ability to bind the same ligand with the active ones or (5) act as protein interaction domains through these their binding interface and so on (Goldberg & Sreelatha, 2023). In plants, studies on the catalytic regulation ability of pseudo-enzymes are rare and still new. (Novikova et al., 2021) reported that the non-enzyme PDX1.2 can positively regulate vitamin B6 production by binding to its active catalytic homologs PDX1.3. Taken together, should OsGDPD13 have weak catalytic activity, it could play the same role as other GDPDs in lipid remodeling, and its silencing will probably be masked by more active GDPDs or functional redundancy by its closest homolog, OsGDPD7, located in chromosome 12. Mehra and Giri (2016) reported that OsGDPD7 upregulated approximately 38-fold after 15 days of P deficiency. On the other hand, non-enzyme activity OsGDPD13 may play a role as a scaffold or allosteric activator/inhibitor of other GDPD enzymes instead, as some bacterial GDPDs are known to form dimer to hexamer higher-order structures (Yip et al., 2011) or obscure GDPD substrates and prevent its hydrolysis by other enzymes, or modulate the functions of other GDPDs. Therefore, *osgdpd13* knock-out or inactivation can account for the change in overall lipid synthesis we have detected.

Besides, interestingly, our results showed that GDPD13 targets many subcellular organelles, including plasma membrane (PM), speckled cytoplasmic, and plasmodesmata (PD). This suggests that GDPD13 may engage the lipid biosynthesis at multiple specific localizations, as do most enzymes involved in lipid biosynthesis metabolism in response to phosphate deficiency. Theoretically, glycerolipids, the main composition of membrane lipids, are synthesized at specific locations such as ER and plastid in the cell and are then distributed to PM and other compartments. Regardless of initially synthesized localization, membranous glycerolipids are generated from phosphatidic acid (PAs) produced directly from PC by phospholipase D or G-3-P and fatty acid *via* the *de novo* lipid synthesis pathway (Kennedy pathway). PAs are further incorporated in DAG or DAG-derivate, which are precursors for glycolipid synthesis in plastid or phospholipid synthesis in the ER (Michaud & Jouhet, 2019; Reszczyńska & Hanaka, 2020; Verma et al., 2021; Yu et al., 2021). By combining with Lipid Acyl Hydrolase (LAH), the GDPD family is involved in the hydrolysis of phospholipids to form G-3-P, which is the primary precursor for PA synthesis *via de novo* lipid biosynthesis pathways (Zhu et al., 2022). Under phosphate-deficient conditions, phospholipids are decomposed and partially replaced by DGDG (Yu et al., 2021). Jouhet and coworkers (2003) have suggested that the DAGs backbone coming from degraded PCs may be used as a substrate for DGDG synthesis in plastids in response to Pi deficiency due to the similarity between FAs composition from decomposed PCs and DGDG molecules. Thus, plasma membrane– associated PCs need to be degraded, and the resulting PC-derivate, PA, and DAG can be trafficked to plastid for DGDG synthesis (Michaud & Jouhet, 2019). Previously, diverse subcellular locations of plant lipid–hydrolyzing protein induced by Pi starvation suggested that the degradation of the phospholipid head group can take place in PM and cytosolic fractions. The membrane–associated non-specific phosphate lipase C AtNPC4 (Nakamura et al., 2005), the vacuoles-mitochondria and vacuoles-plastids membrane contact sites phospholipase D AtPLDζ2 (Yamaryo et al., 2008), cytosolic phospholipase C AtNPC5 (Gaude et al., 2008) the plastidic LPTD1 (Hsueh et al., 2017), the cytoplasmic ZmGPX-PDE1-GFP (J. Wang et al., 2021), the CaGDPD1 targeting ER and other endomembranes (Mehra & Giri, 2016), were reported to hydrolysis of phospholipid backbones from PM and organelle membrane in response to low Pi stress. Moreover, it should be noted that some phospholipid and phospholipid–derived hydrolyzing enzymes change substrate preference according to their subcellular localization. PI-PLC protein families, along with GDPD, are classified together as PLC-like phosphodiesterases by the SCOP database, as an example. Plant PI-PLCs, such as soybean PI-PLC, AtPLC4, and *Petunia* PI-PLC, were detected in both PM in the cytosol. Although most of the plant PI-PLCs can hydrolyze phosphatidylinositol (PI) and PtdIns(4,5)P2, membrane–associated and cytosolic proteins differ in their substrate preference. While cytosolic PI-PLC requires a millimolar mount of Ca2+ for its preferred substrate, phosphatidylinositol (PI), the membrane-associated need merely micromolar amounts of calcium to hydrolysis its priority substrate PtdIns(4,5)P2 (Rupwate & Rajasekharan, 2012).

In addition, it is worth noting that OsGDPD13 targets PD, one type of membrane contact sites connecting the PM and ER. PM and ER are compartments directly involved in lipids biosynthesis and trafficking in plant cells. The ER is the primary place for phospholipids synthesis. The ER extends throughout the cell and communicates extensively with PM and other organelles via membrane contact sites. Among them, PD serves as a multifunctional platform for the non-vesicular movement of lipids, ions, and many different signaling molecules; protein enzymes, in particular the phosphatases, catalyze substrates in the regulatory network of many cellular processes such as lipid metabolism (Li et al., 2021).

Taken together, all of the above shows that OsGDPD13 was involved in the regulation of root development and lipid membrane remodeling under Pi deficiency conditions. Although *in silico* predictions of putative proteins indicated that OsGPDG13 has weak activity to GPDG substrates, we propose that OsGDPD13 affects the enzymatic activity of other GDPDs in some unproven direct or indirect way, thereby altering the G-3-P pool or PA accumulation as well as degrades partly phospholipids.

## CONCLUSION

In this study, we discover the absence of the *OsGDPD13* gene in the *Indica* rice reference genome. The knockout mutant lines of *OsGDPD13* genes have been successfully generated. The phenotypic analysis of mutant lines confirmed the abnormal phenotype compared to wild-type Kitaake regarding the number of crown roots, shoot length, and phospholipids profile under Pi-starvation treatment. This new data confirms the involvement of *OsGDPD13* rice root development and the phospholipid remodeling under Pi deficient conditions. These results provide new information for an in-depth understanding of the molecular function of the *OsGDPD13*, contributing to the improvement of crop plants with higher PUE.

## LIST OF ABBREVIATIONS

CAREs: Cis-acting regulatory elements
GLs: Glycolipids
PLs: Phospholipids
FA(ME): Fatty acid (methyl ester)
TLC: Thin-layer chromatography
GC-MS: Gas chromatography - Mass spectrometry
GDPD: Glycerophosphodiester phosphodiesterase
GPC: Glycerophosphocholine
GPE: Glycerophosphoethanolamine
GPI: Glycerophosphoinositol
GPG: Glycerophosphoglycerol
PAH: Phosphatidic acid phosphohydrolase
PE: Phosphatidylethanolamine
PC: Phosphatidylcholine
PG: Phosphatidylglycerol
PI: Phosphatidylinositol
MGDG: Monogalactosyldiacylglycerol
DGDG: Digalactosyldiacylglycerol
SQDG: Sulfoquinovosyldiacylglycerol
PSRs: Phosphate stress/starvation responses
PUE: Pi utilization efficiency
Pi: inorganic phosphorus
WT: wild-type

## Authorship contribution statement

Tuan-Anh Tran, Tien Van Vu, and Jae-Yean Kim designed and constructed the CRISPR/Cas9 system to inactivate GPDG13. Thi Linh Nguyen, Hoang Ha Chu, and Phat Tien Do contribute to generating a collection of mutant rice lines for GDPD13 and phenotyping the transgenic lines under low Pi stress. Tam Thi Thanh Tran worked out WGS and bioinformatics analysis. Anh-Tuan Tran performed the 3D structure and docking studies. Kieu Thi Xuan Vo and Jong-Seong Jeon performed subcellular localization of OsGDPD13. Dang Thi Thuy Duong and Nguyen Phuong Nhue worked out the phenotyping of knockout OsGDPD13 and phospholipid analyses under control and treated conditions and contributed to performing the experiments and data analysis. Huong Thi Mai To designed and directed the project and wrote the manuscript.

## Supplementary 1

**Figure S1.**
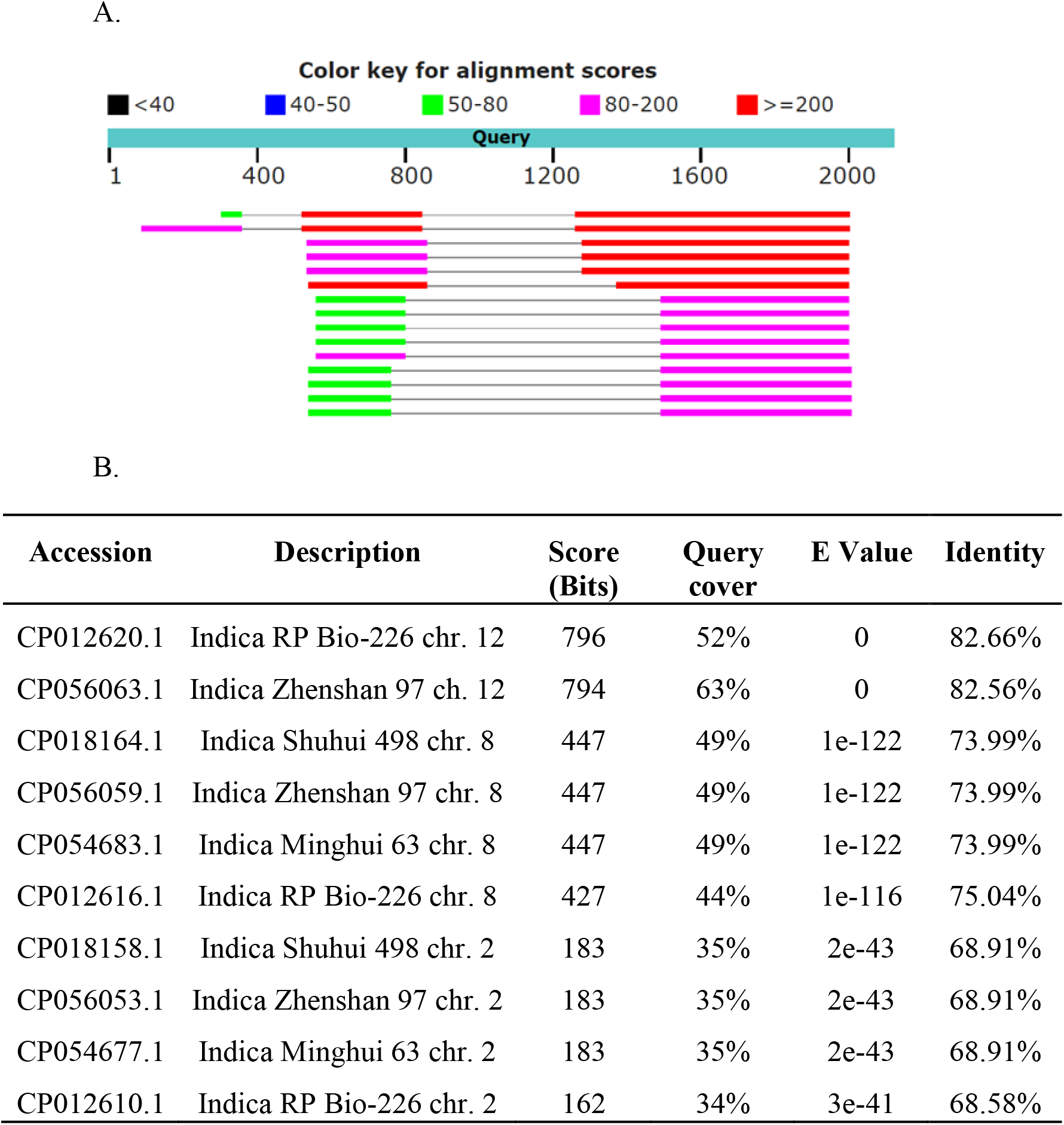
BLASTn homology search of the OsGDPD13 gene against available Indica genomes in the NCBI GenBank database presented as a graphical overview of alignment hits (**A**) and a list of the top 10 best hits (**B**).

## Supplementary 2

### Inheritance and segregation of *OsGDPD13*-induced mutations

The inheritance and segregation of *OsGDPD13* targeted mutations were assessed in the progenies of two representative edited lines #7.2 and #8.1 using the PCR gel electrophoresis and Sanger sequencing. PCR-genotyping analysis of 12 randomly selected T1 progenies of the mutant line #7.2 showed that the two alleles (Figure 3D) of this biallelic mutant were expectedly segregated following Mendelian inheritance patterns with the ratio 1:2:1 (Figure S2A, upper panel). The sequencing data of amplicons from the T1 progenies of line 7.2 were consistent with PCR-gel running results (Figure S2A, bottom panel). In the case of mutant line #8.1, all the examined T1 plants displayed the same single-shifted band as seen in the T0 generation on the agarose electrophoresis (Figure S2B, upper panel). The sequencing data confirmed the homozygous mutation (-7 bp, +1 bp) in T1 progenies of line 8.1 (Figure S2B, bottom panel). Therefore, homozygous mutant alleles of the *OsGDPD13* gene were identified at the T1 generation and used for the T2 analysis.

**Figure S2.**
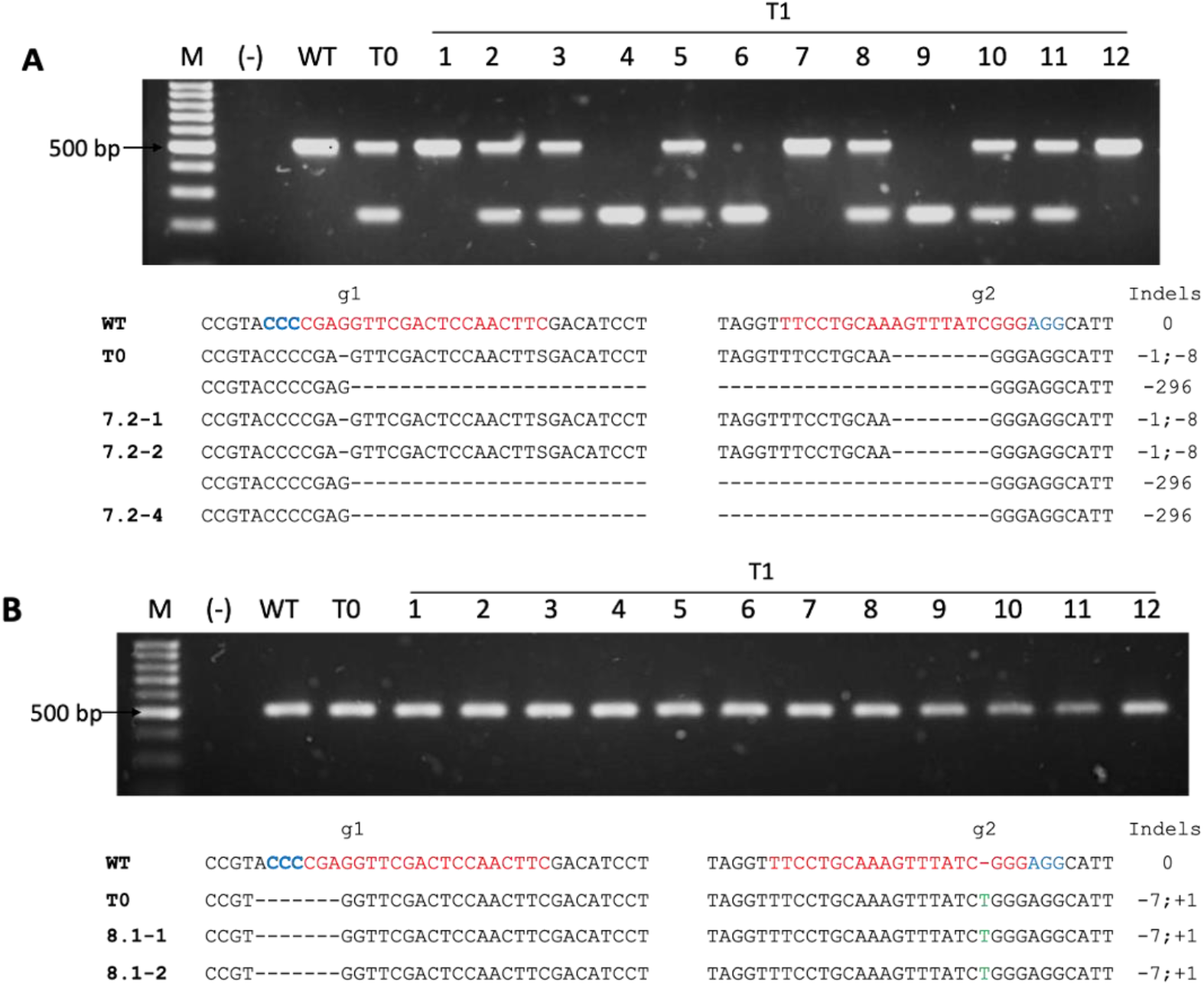
Inheritance and characterization of the *OsGDPD13* targeted mutations at the T1 generation. A. Segregation and inheritance of *osgdpd13* mutants in T1 progenies of the mutant line 7.2. B. Segregation and inheritance of *osgdpd13* mutants in T1 progenies of the mutant line 8.1

**Figure S3.**
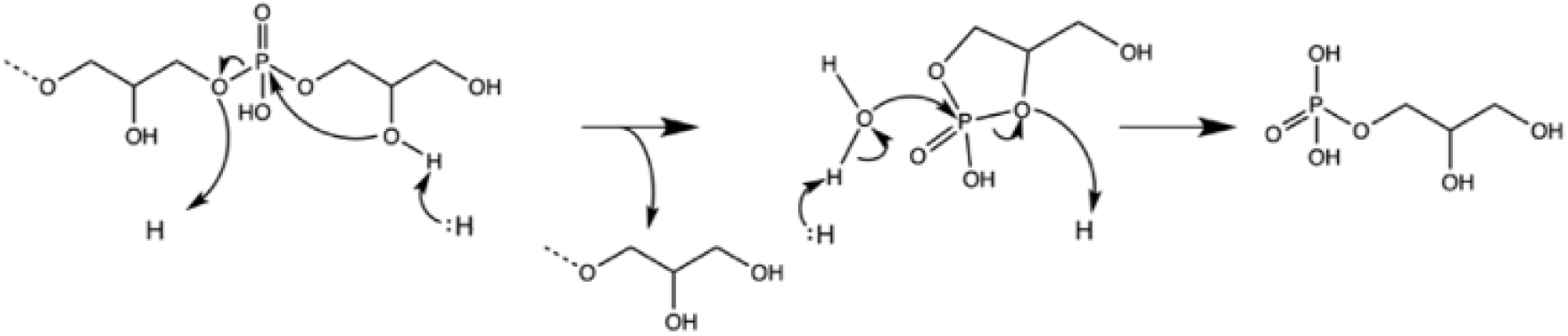
Proposed catalytic mechanism of GDPDs.

**Figure S4.**
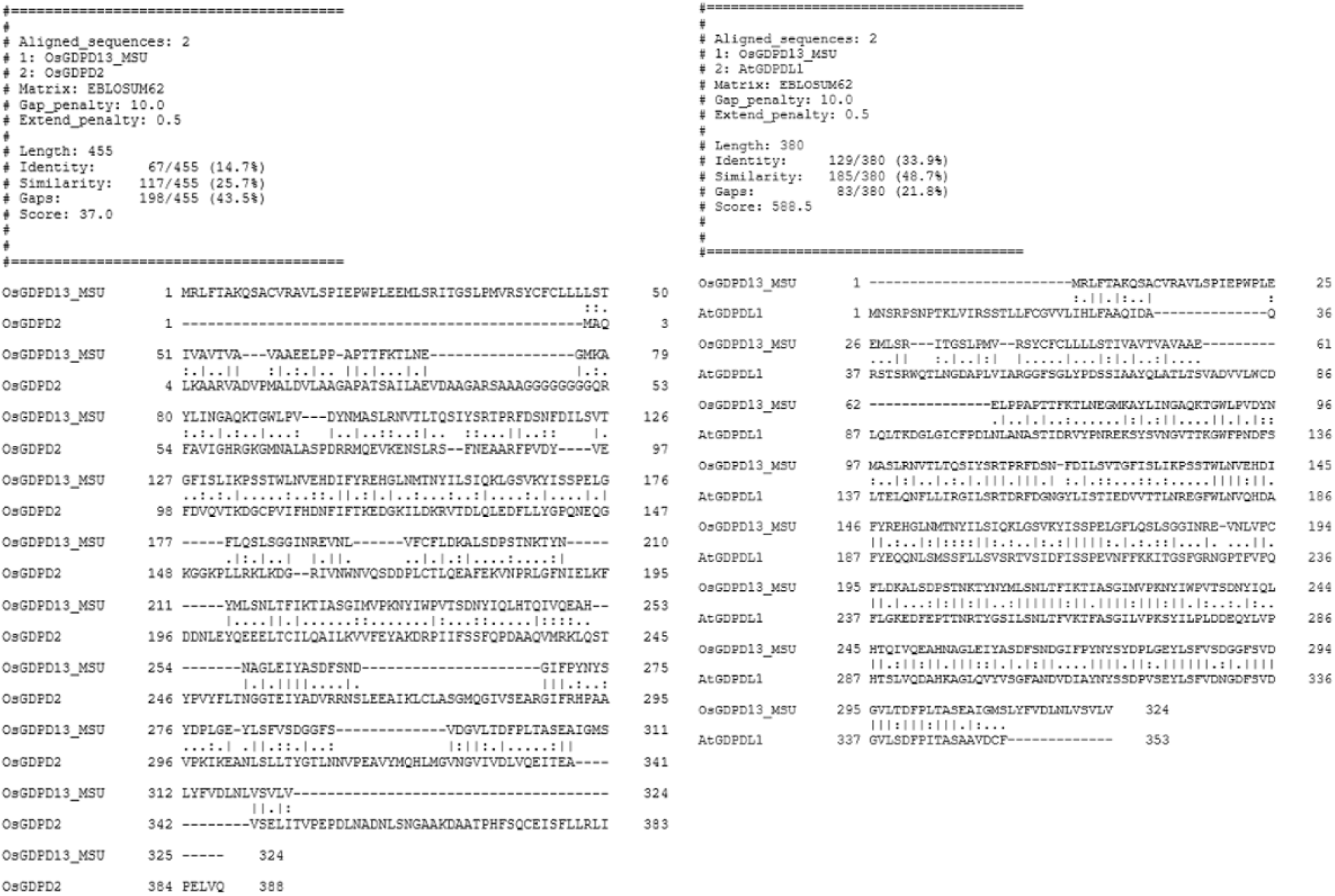
Pairwise alignment of OsGDPD13 (MSU) v. OsGDPD2 and AtGDPDL1 by Clustal Omega, with identity and similarity scores displayed

**Figure S5.**
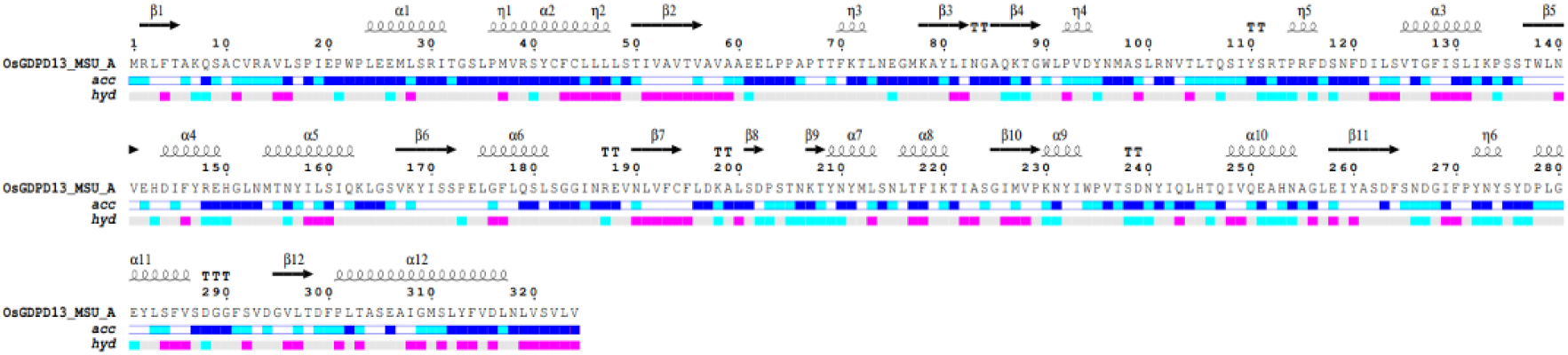
Secondary structures of OsGDPD13 from the MSU database. α-helices are shown as squiggles, β-strands as arrows, 310 helices are η swirls, and TT(T) as strict β- and α-turns, numbered from N- to C-terminal. Below are accompanied solvent accessibility (blue is exposed, cyan is intermediate, white is buried) and hydropathy scales (pink to cyan goes from hydrophobic to hydrophilic)

**Figure S6.**
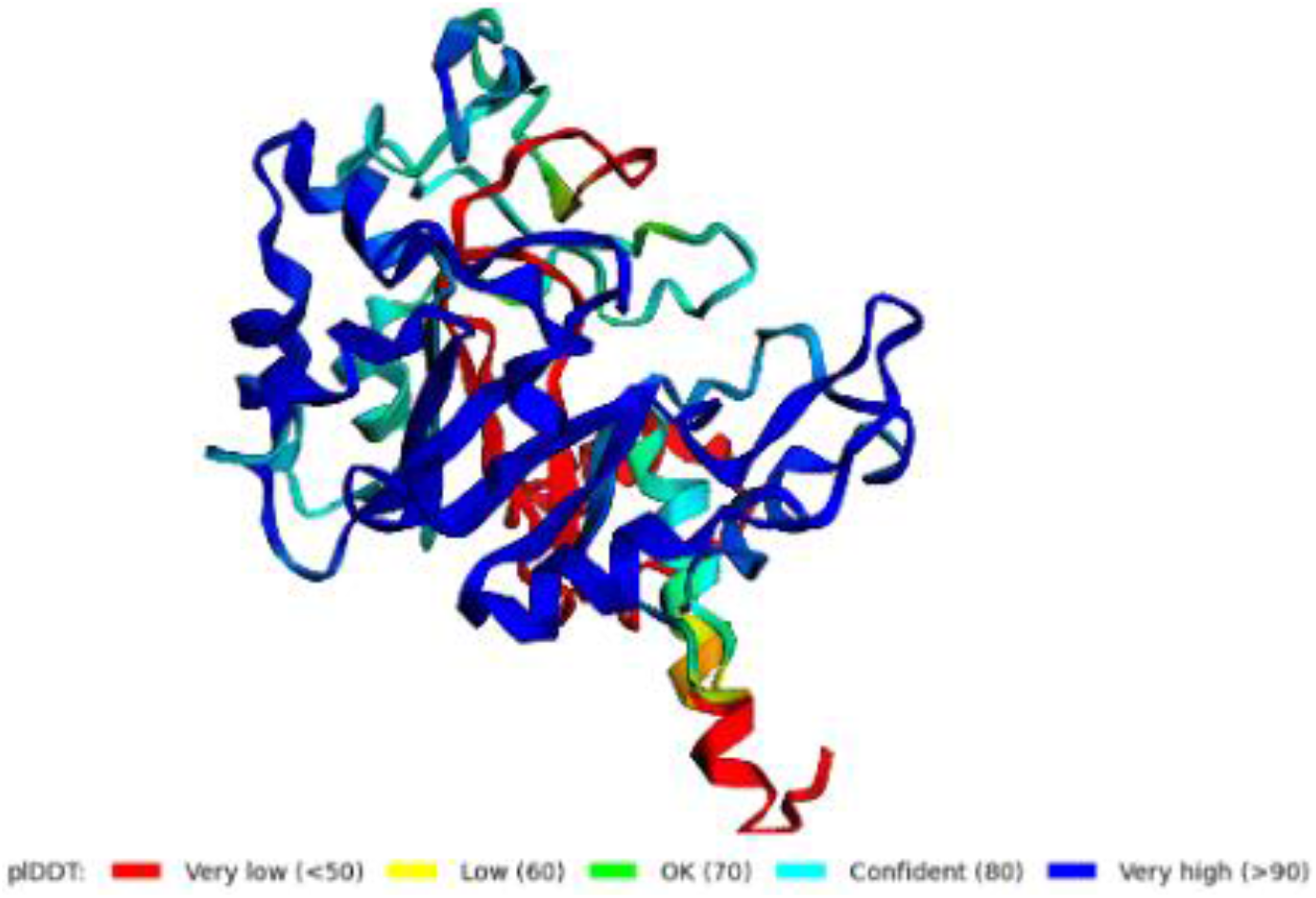
Confidence level of AF2 prediction by pIDDT score [0-100]

**Table S1.**
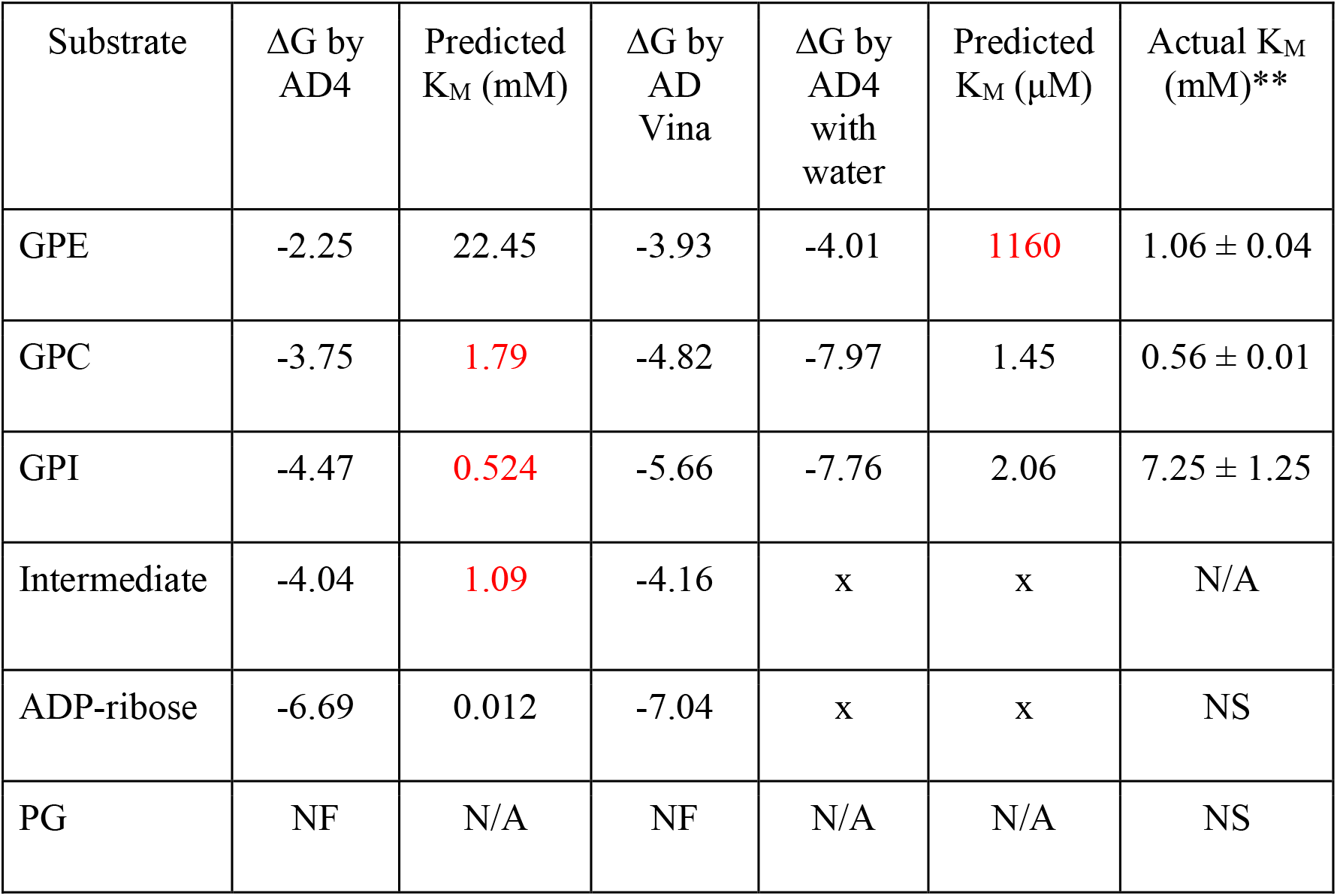
A comparison between binding energy and KM prediction of AD Vina, AD4 and AD4 with hydrated docking on OsGDPD2. Numbers (in red) are within the 0.1-10 μM range

**Table S2.**
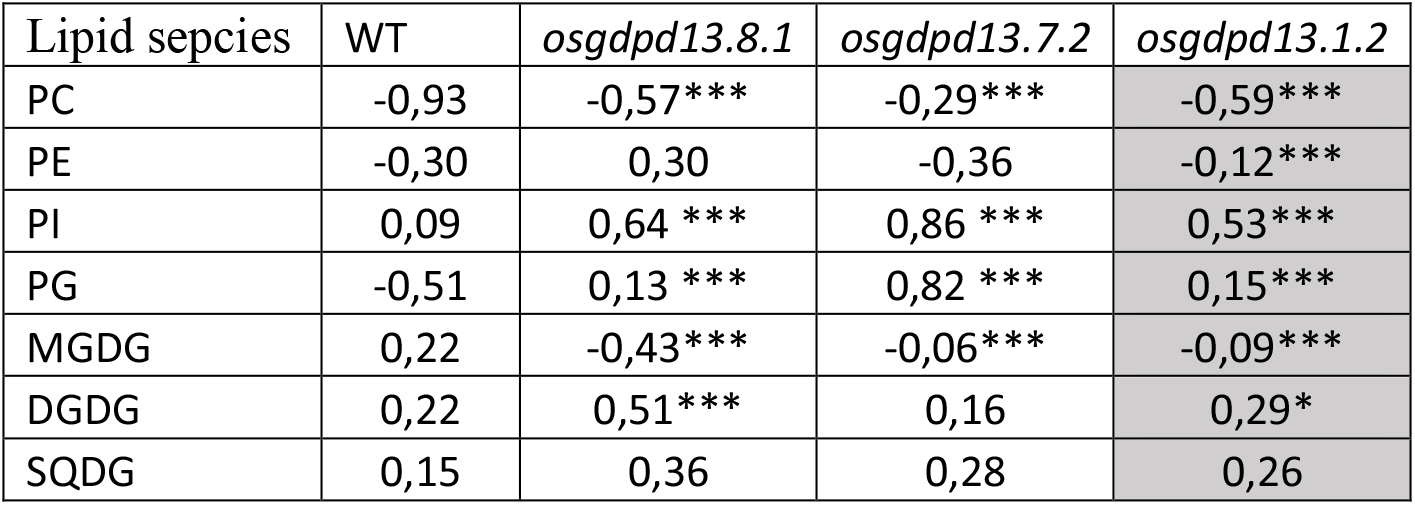
Plasticity index of relative content of lipid species extracted from 21 days – old rice leaves of *osgdpd13* knockout mutant and WT grown under control and low Pi treatment.

| Lipid sepcies | WT | *osgdpd13.8.1* | *osgdpd13.7.2* | *osgdpd13.1.2* |
| --- | --- | --- | --- | --- |
| PC | -0,93 | -0,57*** | -0,29*** | -0,59*** |
| PE | -0,30 | 0,30 | -0,36 | -0,12*** |
| PI | 0,09 | 0,64 *** | 0,86 *** | 0,53*** |
| PG | -0,51 | 0,13 *** | 0,82 *** | 0,15*** |
| MGDG | 0,22 | -0,43*** | -0,06*** | -0,09*** |
| DGDG | 0,22 | 0,51*** | 0,16 | 0,29* |
| SQDG | 0,15 | 0,36 | 0,28 | 0,26 |

